# Admixed populations in the neighbor-joining algorithm: a geometric analysis with five taxa

**DOI:** 10.1101/2024.10.18.619141

**Authors:** Joy Z. Zhang, Wai Tung ‘Jack’ Lo, Michael Stillman, Jaehee Kim

## Abstract

The Neighbor-Joining (NJ) algorithm is a widely used method for constructing phylogenetic trees from genetic distances. While NJ is known to perform well with tree-like data, its behavior under admixture remains understudied. In this work, we present a geometric framework for analyzing the NJ algorithm under a linear admixture model. We focus on three key properties related to clustering order, distance, and topological path length in the resulting NJ trees involving five taxa. Our approach leverages polyhedral geometry to define NJ cones, which correspond to distinct cherry-picking orders and partition the space of dissimilarity vectors. We project dissimilarity vectors with admixture into a lower-dimensional space without admixture, defining polyhedral regions induced by NJ cones that satisfy specified properties. We compute the exact probabilities that these properties hold by directly calculating the volumes of the induced NJ cones and compare them with Monte Carlo integration and standard NJ simulation methods. Our results show that the property on clustering order is always satisfied, while the other properties are highly probable but depend on the admixture fraction. We also prove that certain induced NJ cones have zero volume, indicating that the corresponding NJ tree topologies are infeasible under admixture. We have implemented our methods as a publicly available module `NeighborJoining` within `Macaulay2`, providing an efficient tool for analyzing NJ cones and their properties. This work provides new insights into the geometric structure inherent to the NJ algorithm in the presence of admixture, identifying the conditions under which admixture influences the resulting phylogenetic trees.

## 1 Introduction

The Neighbor-Joining (NJ) algorithm [1] is a widely used distance-based method for inferring phylogenetic trees. Given a pairwise distance matrix, or equivalently a dissimilarity vector, NJ constructs an unrooted binary tree by iteratively merging pairs of nodes according to a specific criterion. When the input distances are additive [2] or nearly additive [3], NJ accurately reconstructs the underlying true tree [4, 5]. However, distances derived from empirical data often deviate from additivity, especially in cases of non-tree-like evolution, such as admixture events resulting from recent gene flow between distinct source populations. Admixed populations, which result from recent mixtures of distinct source populations, often exhibit unique behaviors in NJ trees. Empirical studies across diverse species and genetic datasets [6–12] have demonstrated that an admixed population appears as a short branch along the path connecting its two source populations in an inferred NJ tree. However, the rigorous theoretical understanding of these observed behaviors, including the exact probabilities and the specific conditions under which such patterns arise in NJ trees, remains poorly understood.

Under two-way linear admixture, Kopelman et al. [13] formally defined three key properties that characterize the clustering order, branch lengths, and topological structure of the NJ tree in the presence of admixture: (1) *antecedence of clustering*, where the admixed population clusters with one of its source populations before the two source populations cluster; (2) *intermediacy of distances*, where the distance between the admixed population and each source population is less than the distance between the two source populations; and (3) *intermediacy of topological path lengths*, where the number of edges between the admixed population and each source population is less than or equal to the number of edges separating the two source populations. Kim et al. [14] further investigated these properties through systematic simulations, estimating the approximate probabilities that a random admixed dissimilarity vector satisfies each property. They concluded that, although these properties can be violated, the presence of admixture leads to the properties being satisfied more frequently than expected by chance.

In this work, we introduce a formal geometric framework for studying the behavior of the NJ algorithm under a two-way linear admixture model, with a special focus on the case of five taxa. Unlike previous approaches that rely on empirical or simulation-based methods, our framework enables exact computation of the probabilities that NJ trees satisfy specified properties involving admixture. Our method leverages NJ cones [15, 16], each corresponding to a distinct cherry-picking order, partitioning the space of dissimilarity vectors. By projecting admixed dissimilarity vectors onto a lower-dimensional space without admixture, we define the polyhedral structure induced by the NJ cones. The volumes of these induced NJ cones provide a direct measure of the probability that a random admixed dissimilarity vector satisfies the three specified properties. We have implemented our computational framework as a module, `NeighborJoining`, within `Macaulay2` [17] for public use. Our geometric approach extends the theoretical understanding of the NJ algorithm, offering deeper insights into the structural constraints on the NJ trees imposed by admixture.

## 2 Background and definitions

### 2.1 Neighbor-joining algorithm

The NJ algorithm [1] is an iterative procedure for constructing a phylogenetic tree from an input dissimilarity map of samples. At each iteration of the NJ algorithm, a pair of nodes is selected and merged into a new internal node, which then replaces the original pair. This iterative process continues until an unrooted binary tree, with the samples as tips, is fully constructed. In this work, we restrict our attention to unrooted labeled binary trees, hereafter referred to simply as trees.

The pair selection process (“cherry picking”) in the NJ algorithm corresponds to the formation of half-spaces, which define NJ cones [15]. Each distinct cherry-picking sequence corresponds to an NJ cone. As a result, the output of the NJ algorithm—a specific cherry-picking order for each input dissimilarity map—partitions the space of dissimilarity maps into their respective NJ cones. This section provides an overview of the NJ algorithm within the framework of polyhedral geometry.

#### 2.1.1 Dissimilarity map

The NJ algorithm, as a distance-based method, requires a dissimilarity matrix as input. Each entry in this matrix represents the pairwise dissimilarity between samples, defined by a dissimilarity map on the sample space. This map satisfies the properties of non-negativity, identity, and symmetry, while potentially relaxing the triangle inequality, thus operating as a semi-metric.

##### Definition 1 (Dissimilarity matrix and dissimilarity vector)

We denote the total number of initial samples by *N* and let *n* ∈ [*N*] represent the number of taxa remaining at a given iteration. Here, [*N*] denotes the set {1, …, *N* } for a positive integer *N* . The index set corresponding to these *n* taxa is denoted by *I*^(*n*)^, where *n* = |*I*^(*n*)^|. The *dissimilarity matrix* **D**_*n*_ for *n* taxa is an *n* × *n* symmetric matrix with a zero diagonal, where each off-diagonal entry represents the dissimilarity between a pair of taxa.

We define the *dissimilarity vector* **d**^(*n*)^ for *n* taxa as a column vector of length 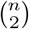, composed of the lower-triangular entries of the dissimilarity matrix **D**_*n*_, arranged in lexicographical order based on the taxon indices. The entry in the *i*-th row and *j*-th column (*i > j*) of **D**_*n*_ is bijectively mapped to the 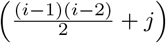 -th position in **d**^(*n*)^ [16].

For example, when *n* = 5 with *I*^(*n*)^ = [5],

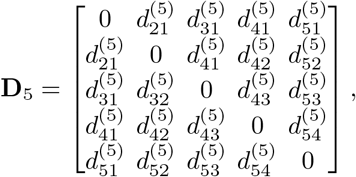

and

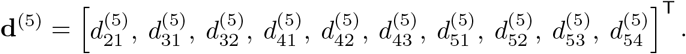

#### 2.1.2 The Q-criterion

At each step of the NJ algorithm, a pair of taxa is selected based on the Q-criterion, a linear transformation mapping each entry 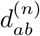 of the dissimilarity vector to its corresponding Q-value *q*_*ab*_, as defined below.

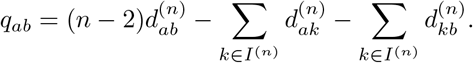

This linear transformation can be represented by a 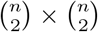 matrix, which we refer to as the A-matrix.

##### Definition 2 (A-matrix)

Let the indices *i* and *j* correspond to the pairs of taxa (*a, b*) and (*c, d*), respectively. The *A-matrix* **A**^(*n*)^ is defined as a 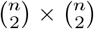 matrix, where each entry 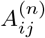 is given by:

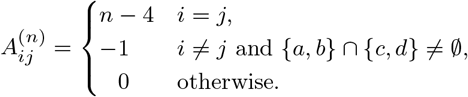

For example, when *n* = 5,

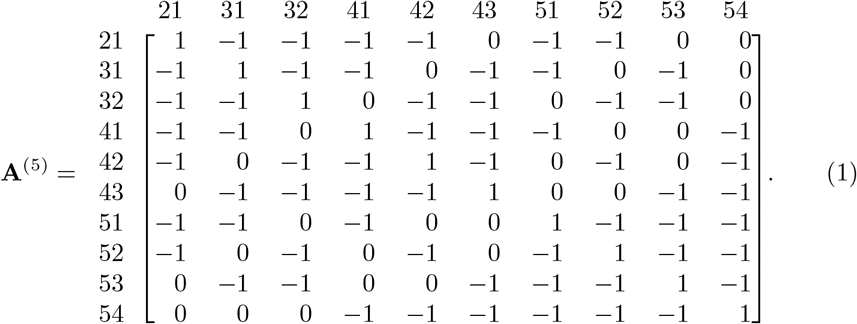

We define the Q-vector **q**^(*n*)^ as **q**^(*n*)^ = **A**^(*n*)^**d**^(*n*)^. The pair of taxa to be merged in the current iteration corresponds to the index of the Q-vector with the minimum value: 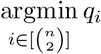.

#### 2.1.3 Updating the dissimilarity vector and tree construction

After selecting a pair of taxa for merging, a new node representing the pair is created. The newly created nodes are indexed as *N* + 1, *N* + 2, …, 2*N* − 2, with node *N* + *i* introduced in the *i*-th iteration. The original pair is removed from the set of taxa in the subsequent iteration, decreasing the number of taxa to be processed by one. The dissimilarity vector **d**^(*n*)^ is then updated to **d**^(*n*−1)^ by removing the entries associated with the original pair and introducing new entries that represent the dissimilarities between the newly created node and all remaining taxa.

Formally, let *a, b* ∈ *I*^(*n*)^ be the pair of taxa selected at a given step, with *u* representing the newly created node for the pair. Then, *I*^(*n*−1)^ = {*u*} ∪ *I*^(*n*)^ \ {*a, b*}, and the entries of the updated dissimilarity vector **d**^(*n*−1)^ are defined as follows:

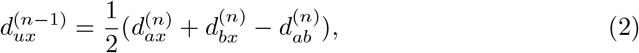

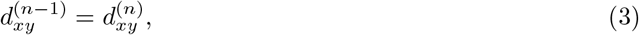

where *x, y* ∈ *I*^(*n*)^ \ { *a, b*} . This process of updating the dissimilarity vector at each step of the algorithm can be formalized as a linear transformation [16].

##### Definition 3 (R-matrix)

We define the *R-matrix* as a 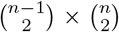 matrix **R**^(*n*)^, such that **d**^(*n*−1)^ = **R**^(*n*)^**d**^(*n*)^. Let *k*_*cd*_ and *k*_*ef*_ denote the positions of the pairs (*c, d*) and (*e, f*), respectively, in the lexicographically ordered set {(*x, y*) | *x, y* ∈ [5], *x > y*}. The entry 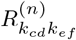 in the *k*_*cd*_ row and the *k*_*ef*_ column is defined as:

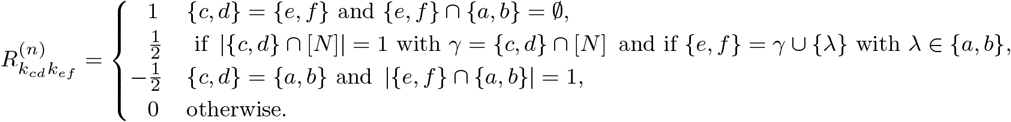

For example, if we start with *n* = 5, and taxa 2 and 3 are selected in the first iteration, the resulting dissimilarity vector **d**^(4)^ is:

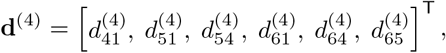

and the matrix **R**^(5)^ that updates the dissimilarity vector from **d**^(5)^ to **d**^(4)^ is given by:

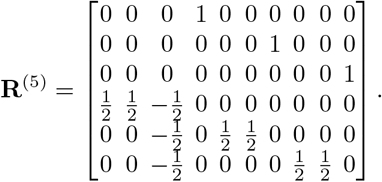

After merging taxa *a* and *b* into a newly created node *u*, the node pairs (*a, u*) and (*b, u*) define edges *e*_*au*_ and *e*_*bu*_, respectively, on the inferred NJ tree, with their lengths *β*_*au*_ and *β*_*bu*_ given by:

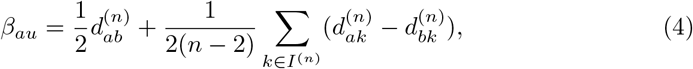

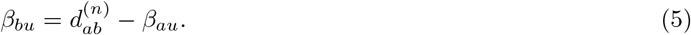

The final NJ tree with *N* leaves is obtained after *N* − 2 iterations. To quantify distances within this tree, we introduce the tree metric (or additive metric) [2, 5] that defines the pairwise distances between the leaves.

##### Definition 4 (Minimum path length between nodes)

Let *V* = [2*N* − 2] denote the set of nodes in the final NJ tree. We define the function *𝓁* : *V* × *V* → ℝ_≥0_ as the map that assigns to each pair of nodes in the tree the *minimum path length* between them. Specifically, for any two nodes *i, j* ∈ *V*, *𝓁*(*i, j*)—denoted by *𝓁*_*ij*_ —represents the sum of edge lengths along the shortest path connecting *i* and *j* within the tree. If *i* and *j* are adjacent, *𝓁*_*ij*_ is precisely *β*_*ij*_, the length of the edge directly connecting them.

##### Definition 5 (Tree metric)

By restricting *𝓁* to the set of leaves [*N*], we define the *tree metric δ* as a column vector of length 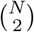, where each entry *δ*_*ij*_ corresponds to the minimum path length *𝓁*_*ij*_ between leaves *i, j* ∈ [*N*] of the tree.

#### 2.1.4 NJ cones

In Sections 2.1.2 and 2.1.3, we formalized the NJ algorithm as a sequence of linear transformations applied to the input dissimilarity vector, with the cherry-picking order determined by the Q-criterion, represented as a set of linear inequalities at each iteration. This process defines a geometric structure known as NJ cones [15], where each cone corresponds to a unique NJ tree, thereby partitioning the space of input dissimilarity vectors. This section provides a mathematical description of this structure.

Let *a* and *b* be the pair of taxa selected at a given step with *n* taxa remaining, and let 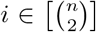 denote the index corresponding to the position of the pair (*a, b*) within **d**^(*n*)^. Denote by 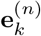 the *k*-th standard basis column vector in 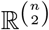, with a 1 in the *k*-th position and 0 in all other positions. Then,

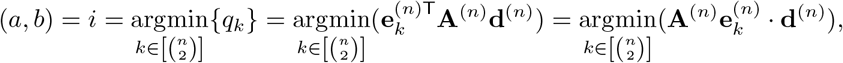

where the final step follows directly from the symmetry of the A-matrix.

Since *i* corresponds to the index of the minimum Q-criterion value, the following inequality holds for all 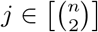:

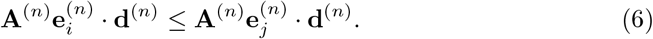

For given *i*, and for each *j* ∈ *I*^(*n*)^, Eq. 6 defines a half-space in the space of input dissimilarity vectors 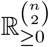:

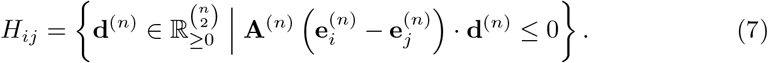

If an input dissimilarity vector **d**^(*n*)^ satisfies the inequality in Eq. 7, it lies within the corresponding half-space *H*_*ij*_. When *i* = *j*, the inequality holds trivially, resulting in 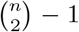 non-trivial half-spaces for each cherry-picking among *n* nodes. Thus, the *i*-th pair is selected if and only if **d**^(*n*)^ lies within the intersection of all non-trivial half-spaces generated in that step, formally expressed as

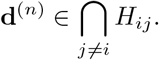

The total number of half-spaces required to reconstruct a tree with *N* samples is obtained by summing the number of half-spaces generated at each step of the algorithm: 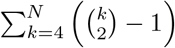. Note that at the iteration where *n* = 3, all entries of the Q-vector are identical because **A**^(3)^ has rank 1 by construction:

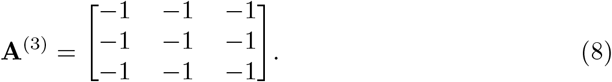

Therefore, only the iterations preceding *n* = 3 contribute to the total set of half-spaces. We now formally define the NJ cone using these half-planes.

##### Definition 6 (NJ cone)

Let 𝒪= (*o*_1_, *o*_2_, …, *o*_*N* −3_) represent a cherry-picking order on a set of *N* taxa, specifying the sequential selection of *N* − 3 cherries, with *o*_*k*_ denoting the *k*-th cherry. At each iteration *k*, define *n*_*k*_ = *N* − *k* + 1 as the number of taxa remaining at the beginning of the iteration, and let 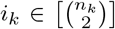 be the index of *o*_*k*_ in the corresponding Q-vector. For each 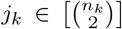 with *j*_*k*_ ≠ *i*_*k*_, define the row vector 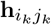 of length 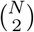, constructed from the comparison of the Q-values 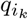 and 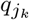 as follows:

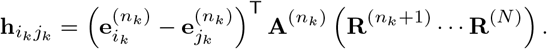

At iteration *k*, there are 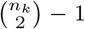 hyperplanes, defined by the vectors 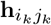, for each 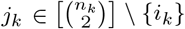. We define the reindexing 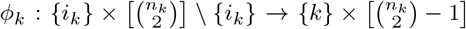, which reassigns the indices *i*_*k*_ and *j*_*k*_ as follows:

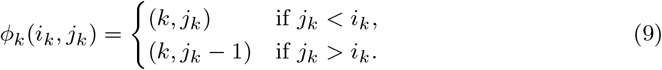

This reindexes each hyperplane 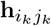 to **h**_*km*_, where 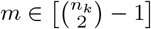.

We define the *NJ cone matrix* 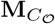 as a 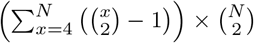 matrix, where rows correspond to hyperplanes associated with the cherry-picking order 𝒪:

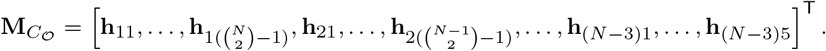

The *NJ cone C*_𝒪_associated wit the cherry-picking order 𝒪is hen defined as:

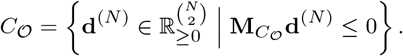

In other words, the NJ cone *C*_𝒪_ represents the region in the space of dissimilarity vectors where the NJ algorithm, following the order 𝒪, produces a unique unrooted labeled binary tree topology, distinguishing reflection-symmetric topologies by assigning them to distinct cones (Table 2).

**Table 1:**
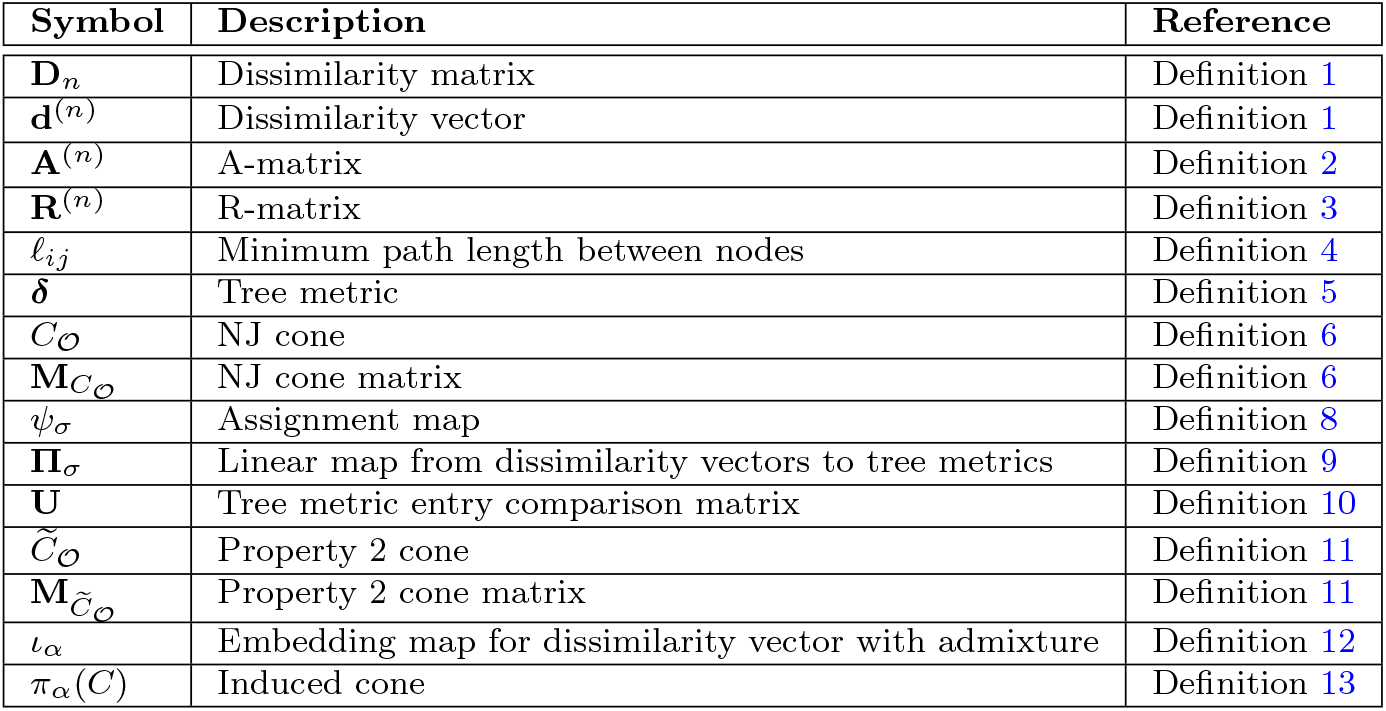
Summary of notations and definitions. This table lists the key symbols and terms used in this work, with references to their formal definitions.

**Table 2:**
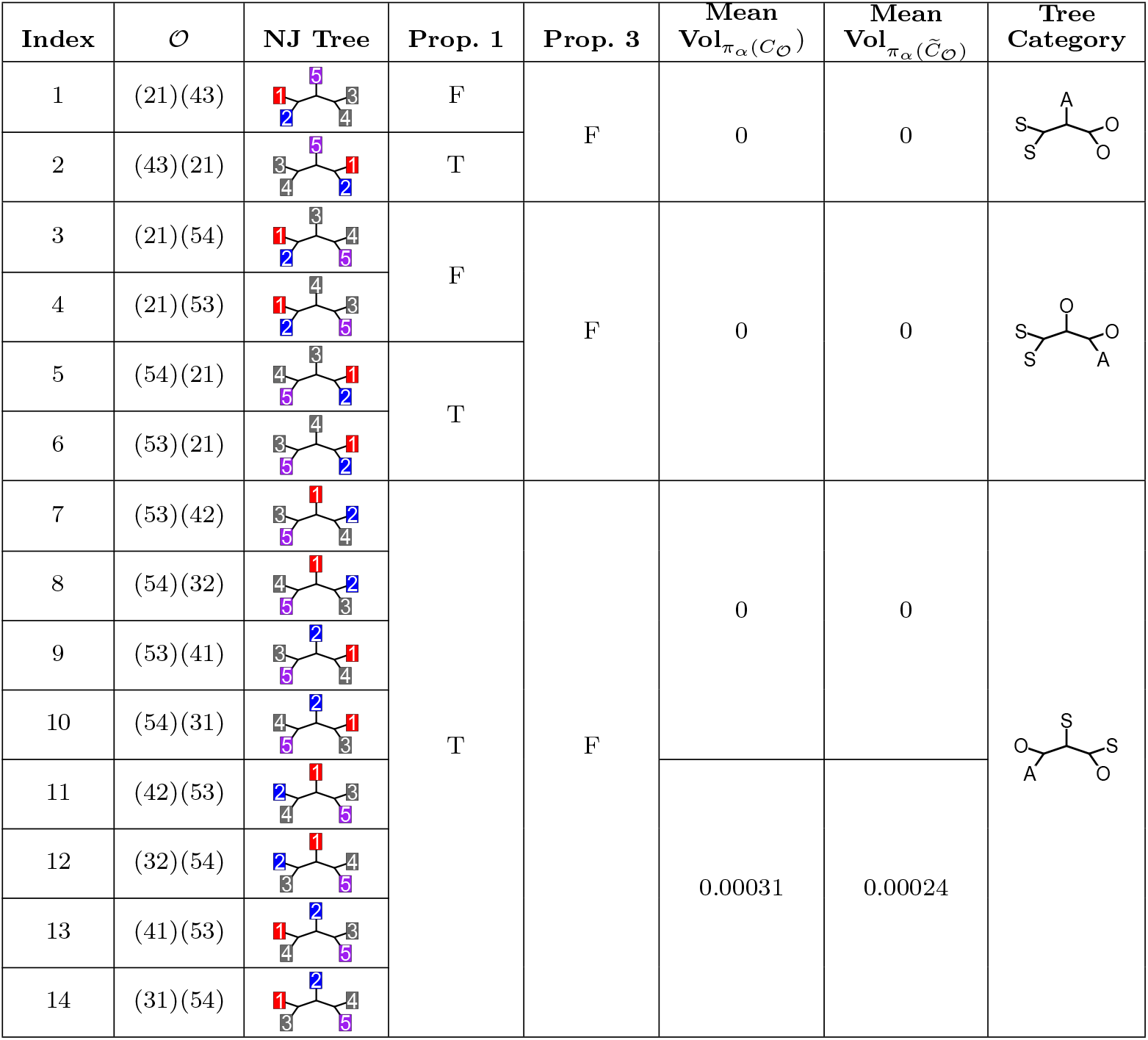
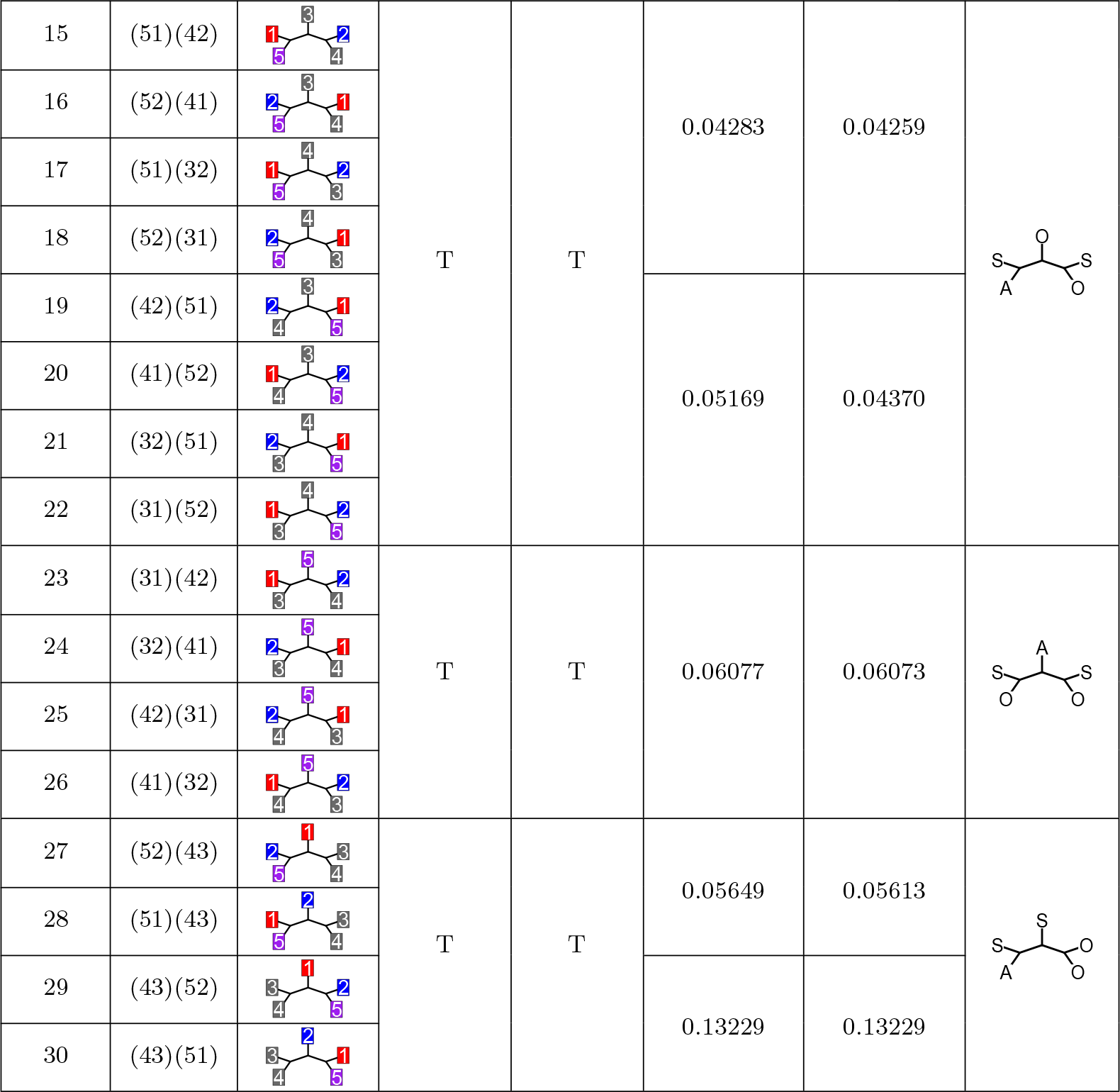
Mean 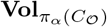 and mean 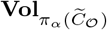 from direct computation. The first column lists the indices of 30 distinct NJ cones *C*_𝒪_ for *N* = 5, while the second column shows the equivalence class of cherry-picking orders for each NJ cone, following the convention in Section 2.1.5. The third column illustrates the labeled tree topology for each order, distinguishing reflection symmetry about the central taxon. The fourth and fifth columns indicate whether the corresponding NJ cone satisfies Property 1 and Property 3, respectively. “T” denotes that the property is satisfied, while “F” indicates it is not. The sixth and seventh columns report the mean volumes of *π*_*α*_(*C*_𝒪_) and 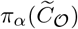, respectively, averaged over 99 values of *α* in {0.01, 0.02, …, 0.99} for each 𝒪 ∈ Ω_all_. These volumes were obtained using the direct computation method (Section 3.3.1). Non-averaged volumes at *α* = 0.01, 0.5, and 0.99 are provided in Table A1. The final column categorizes the 30 NJ cones by tree topology, distinguishing three node types: source taxa (*S*), an admixed taxon (*A*), and other taxa (*O*). This classification yields six distinct labeled tree topologies based on the arrangement of two *S* taxa, one *A* taxon, and two *O* taxa. These topology categories are considered equivalent if they differ only by reflection about the central taxon. Rows are ordered by increasing mean volume within each labeled tree topology category, with NJ cones further sorted by the ascending label of the central taxon.

Each input dissimilarity vector **d**^(*N*)^ either lies strictly within the interior of a single NJ cone or on a boundary shared by multiple NJ cones. The latter occurs if and only if there exists an NJ cone *C*_𝒪_ and a cherry-picking iteration *k* such that, at the *k*-th iteration, a row vector 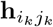 in 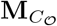 satisfies:

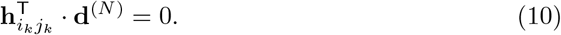

Eq. 10 implies that the entries 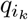 and 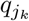 of the Q-vector at the *k*-th iteration are both the minimum among all entries of the Q-vector at that iteration. So the NJ algorithm cannot distinguish picking the cherry *i*_*k*_ or the cherry *j*_*k*_ at the *k*-th iteration. Let *m*_*k*_ denote the number of cherries whose Q-vector entries have the same minimum value at the *k*-th iteration, i.e., there are *m*_*k*_ pairs of nodes satisfying Eq. 10. The cherry-picking order for **d**^(*N*)^ can then select any of these *m*_*k*_ cherries, as their Q-vector entries are all minimum.

Since each iteration determines a single cherry, and there are *N* − 3 iterations that define the half-spaces bounding an NJ cone, the total number of distinct NJ cones containing **d**^(*N*)^ on the boundary is given by multiplying the number of equivalent choices at each iteration, adjusting for the three pairs of identical rows in the A-matrix at the (*N* − 3)-th iteration (Eq. 8):

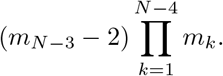

For example, for *N* = 5, if **d**^(5)^ has *m*_1_ = 5 and *m*_2_ = 3, it lies on the boundary shared by (*m*_2_ − 2)*m*_1_ = (3 − 2) × (5 − 0) = 5 distinct NJ cones.

#### 2.1.5 The equivalence relation among cherry-picking orders

While each cherry-picking order belongs to an NJ cone, a single NJ cone can correspond to multiple cherry-picking orders due to the structural properties of the A-matrix (Definition 2). If two rows of **A**^(*n*)^ are identical, the corresponding Q-vector entries are equal for any input dissimilarity vector (Section 2.1.2). In this case, the NJ algorithm arbitrarily selects a cherry from node pairs with the minimum Q-value. This multiplicity induces an equivalence class of cherry-picking orders. The proof of this equivalence relation is straightforward and is therefore omitted.

##### Definition 7 (Equivalence class of cherry-picking orders)

Two cherry-picking orders are *equivalent* if and only if, at every iteration of the NJ algorithm, the corresponding rows of the A-matrix are identical.

For *N* = 5, the set of cherry-picking orders forms thirty distinct equivalence classes, with each class containing six orders [15]. This results from the combinatorial properties of tree topologies under the NJ algorithm. There is one unrooted unlabeled binary tree topology, which can be labeled to form (2*N* − 5)!! = 15 unrooted labeled binary tree topologies. However, the cherry-picking order introduces asymmetry relative to the central edge, thereby doubling the total number of unique NJ cones to thirty. Thus, each NJ cone corresponds to a unique unrooted labeled binary tree topology that accounts for asymmetry with respect to the central node, resulting in a total of thirty distinct NJ cones (Table 2).

The equivalence of six cherry-picking orders per class is due to the ties in the Q-criterion, as defined by the matrices **A**^(4)^ and **A**^(3)^. At the first iteration, the Q-vector is given by **q**^(5)^ = **A**^(5)^**d**^(5)^. Since the rows of **A**^(5)^ are distinct (Eq. 1), no two cherries are guaranteed to have the same Q-value at this step. Assuming, without loss of generality, that nodes 4 and 5 are selected as a cherry in the first step, with node 6 added, the Q-vector at the second step is computed as **q**^(4)^ = **A**^(4)^**d**^(4)^, where **A**^(4)^ is given by:

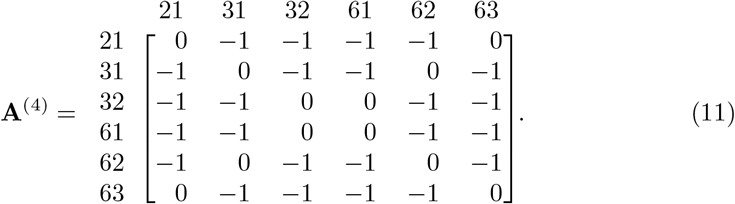

Regardless of the cherry selected in the first step, **A**^(4)^ contains three pairs of identical rows, leading to ties in the Q-values. In the third and final cherry-picking step, **A**^(3)^ has identical rows (Eq. 8), resulting in ties across all node pairs. These ties in both the second and third steps establish the equivalence of six cherry-picking orders.

This equivalence allows for a systematic representation of each NJ cone for *N* = 5 based on the first two cherries. Since each NJ cone corresponds to six equivalent cherry-picking orders, any of these orders can be selected to represent the cone. Among these six, three have their first two cherries composed solely of the original taxa indices {1, …, 5} . We represent the cone by the cherry-picking order where these cherries consist exclusively of the original indices. Let (*a, b*) and (*c, d*) be the first two cherries of an NJ cone, where *a, b, c, d* ∈ [5]. The NJ cone is then denoted by *C*_(*a*,*b*)(*c*,*d*)_. This notation will be used to represent NJ cones with *N* = 5 throughout the remainder of this work.

### 2.2 Populations with admixture

Consider a set of *N* populations, indexed by *I*^(*N*)^ = [*N*] and labeled {*t*_1_, *t*_2_, …, *t*_*N*_ }. Let *t*_*N*_ represent an admixed population from a two-way admixture between source populations *t*_1_ and *t*_2_. Define the admixture fraction *α* ∈ (0, 1) as the proportion of contribution from source population *t*_1_ to the admixed population *t*_*N*_, with the remaining 1 − *α* representing the contribution from the other source population *t*_2_. We employ the linear admixture model as described in Kopelman et al. [13] and Kim et al. [14], where the pairwise genetic distances between the admixed population *t*_*N*_ and other populations are expressed as a linear combination of the distances involving the source populations. For each *i* ∈ [*N*], these distances are given by:

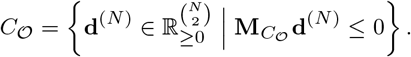

The corresponding dissimilarity matrix **D**^(*N*)^, incorporating the admixed population *t*_*N*_ is given by:

**D**(*N*) =

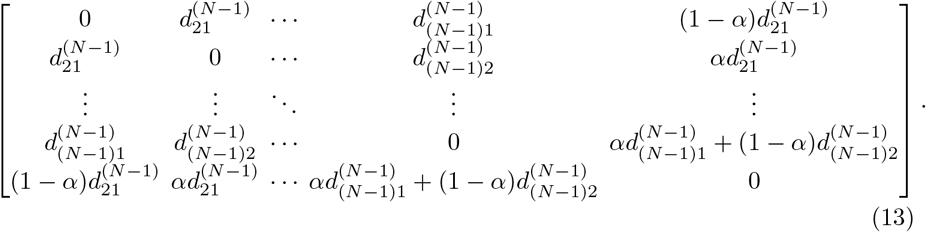

We assume all off-diagonal elements in **D**^(*N*)^ are strictly positive.

#### 2.2.1 Property 1: antecedence of clustering

In the NJ algorithm, Property 1 is satisfied if the admixed taxon clusters with one of the source taxa before the two source taxa are clustered together. Formally, let Γ_*i*_ denote the clade containing taxon *i*, and (Γ_*i*_, Γ_*j*_) represent the agglomeration of two clades. The clustering order satisfies Property 1 in the last two of the following three cases:

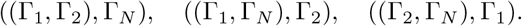

For example, when *N* = 5, Property 1 is satisfied if source taxon 1 and admixed taxon 5 are clustered in the first step, forming a new node 6. In the second step, node 6 clusters with taxon 2, such that the admixed taxon 5 is clustered with source taxon 1 before the source taxa 1 and 2 are joined. Conversely, Property 1 is violated if source taxa 1 and 2 are clustered in the first step.

#### 2.2.2 Property 2: intermediacy of distances

Property 2 states that the distances on the inferred NJ tree (the tree metric; Definition 5) between the admixed taxon and each source taxon are less than or equal to the distance between the two source taxa. Since we are working within the framework of closed polyhedra, the strict inequality “*<*“ from Kopelman et al. [13] is replaced by “ ≤ “. This adjustment does not affect the probability of satisfying or violating the property, since the boundary of a polyhedron has measure zero. Thus, the property requires:

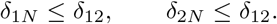

#### 2.2.3 Property 3: intermediacy of topological path lengths

Unlike Property 2, which focuses on branch lengths, Property 3 concerns the topological structure of the NJ tree, which is determined by the cherry-picking order. This property requires that the number of edges along the shortest path between the admixed taxon and each source taxon be less than or equal to the number of edges along the shortest path between the two source taxa. Formally, let *τ*_*ij*_ denote the number of edges on the shortest path between taxa *i* and *j*. Property 3 is then expressed as:

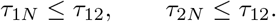

## 3 Methods

Our objective is to compute the probability that a random dissimilarity vector with admixture produces an NJ tree satisfying the three properties defined in Section 2.2. Given that NJ cones partition the space of dissimilarity vectors (Section 2.1.4), associating each NJ cone with these properties enables the identification of dissimilarity vectors that satisfy them. In this section, we present a theoretical framework for computing these probabilities based on the volumes of NJ cones, employing two approaches: (1) Monte Carlo integration and (2) direct calculation. While the methods are demonstrated for *N* = 5, they can be generalizable to cases where *N >* 5.

### 3.1 Classification of NJ cones based on the three properties

Properties 1 and 3 are topological, determined solely by the cherry-picking order associated with the input dissimilarity vector. In contrast, Property 2 is not necessarily satisfied across an entire NJ cone, as it further depends on the branch lengths of the final NJ tree. In this section, we identify the NJ cones whose equivalence classes of cherry-picking orders satisfy Property 1 and, separately, those that satisfy Property 3. We also construct the cones associated with Property 2 using a linear transformation that maps input dissimilarity vectors to the corresponding tree metrics in the final NJ tree.

#### 3.1.1 NJ cones satisfying Property 1 (antecedence of clustering)

We first identify NJ cones satisfying Property 1 for *N* = 5. Property 1 holds if the admixed taxon clusters with either source taxon before the source taxa cluster with each other. Within an equivalence class of an NJ cone, some cherry-picking orders may satisfy Property 1, while others may not. We classify NJ cones into three categories based on the number of cherry-picking orders in their equivalence class that satisfy Property 1. Since ties in the Q-criterion define these equivalence classes, we deem an NJ cone to satisfy Property 1 if at least one cherry-picking order within its equivalence class satisfies it.

##### Type 1: NJ cones whose equivalence class consists entirely of cherry-picking orders that satisfy Property 1

If the first cherry in a cherry-picking order is either (5, 1) or (5, 2), where one taxon is the source and the other the admixed taxon, the corresponding NJ cone satisfies Property 1 and is classified as Type 1. In this case, the source and the admixed taxa form a new node, which is subsequently clustered with the remaining source taxon. Thus, all cherry-picking orders within this equivalence class have this property. For example, the NJ cone corresponding to the NJ tree in Figure 1 is classified as a Type-1 cone. All six cherry-picking orders associated with this NJ cone satisfy Property 1: (5, 1)(4, 2)(7, 6), (5, 1)(4, 2)(7, 3), (5, 1)(4, 2)(6, 3), (5, 1)(6, 3)(7, 4), (5, 1)(6, 3)(7, 2), and (5, 1)(6, 3)(4, 2). The complete set of NJ trees corresponding to Type-1 cones is provided in Figure B1.

**Fig. 1:**
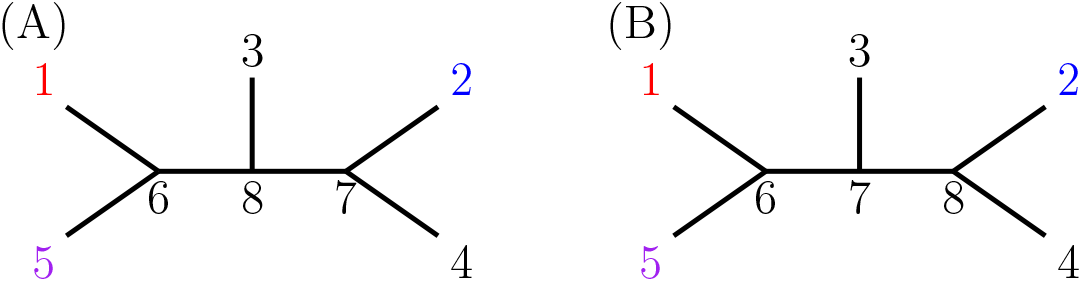
Example NJ tree corresponding to a Type-1 NJ cone satisfying Property. **1**. The two trees shown are the same NJ tree but differ only in their internal node labels, resulting from different cherry-picking orders within the same equivalence class. **(A)** Labeled tree topology, with internal node labeled, from the cherry-picking orders: (5, 1)(4, 2)(7, 6), (5, 1)(4, 2)(7, 3), and (5, 1)(4, 2)(6, 3). **(B)** Labeled tree topology, with internal node labeled, from the cherry-picking orders (5, 1)(6, 3)(7, 4), (5, 1)(6, 3)(7, 2), and (5, 1)(6, 3)(4, 2).

##### Type 2: NJ cones whose equivalence class contains at least one, but not all, cherry-picking orders that violate Property 1

If an equivalence class of an NJ cone includes at least one cherry-picking order that satisfies Property 1, but not all, we classify the NJ cone as Type 2 and satisfying Property 1. This convention is adopted because the equivalence class arises from ties in the Q-criterion, where any tied pairs can be selected randomly. Thus, if any cherry-picking order within the equivalence class satisfies Property 1, we designate the entire NJ cone as satisfying Property 1. For example, the NJ cone corresponding to the NJ tree in Figure 2 is a Type-2 cone. The cherry-picking order, (3, 1)(4, 2)(7, 6) violates Property 1, whereas the other cherry-picking orders in the same equivalence class—(3, 1)(4, 2)(6, 5), (3, 1)(4, 2)(7, 5), (3, 1)(6, 5)(7, 4), (3, 1)(6, 5)(7, 2), and (3, 1)(6, 5)(4, 2)—satisfy it. The complete set of NJ trees corresponding to Type-2 cones is listed in Figure B2.

**Fig. 2:**
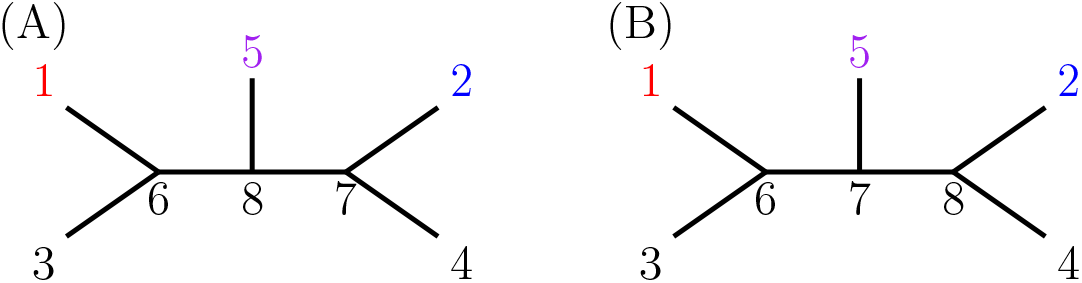
Example NJ tree corresponding to a Type-2 NJ cone satisfying Property. 1. The two trees shown are the same NJ tree but differ only in their internal node labels, resulting from different cherry-picking orders within the same equivalence class. **(A)** Labeled tree topology, with internal node labeled, from the cherry-picking orders: (3, 1)(4, 2)(7, 6), (3, 1)(4, 2)(6, 5), and (3, 1)(4, 2)(7, 5). **(B)** Labeled tree topology, with internal node labeled, from the cherry-picking orders: (3, 1)(6, 5)(7, 4), (3, 1)(6, 5)(7, 2), and (3, 1)(6, 5)(4, 2).

##### Type 3: NJ cones whose equivalence class consists entirely of cherry-picking orders that violate Property 1

If the first cherry in a cherry-picking order consists of the source taxa, (2, 1), the corresponding NJ cone is classified as Type 3 and necessarily violates Property 1. This violation occurs because all cherry-picking orders in this equivalence class start with (1, 2) as the first cherry, and the admixed taxon 5 can only join (2, 1) after these two source taxa have already been clustered. For example, the NJ cone corresponding to the NJ tree in Figure 3 is classified as a Type-3 cone. All six cherry-picking orders associated with this NJ cone violate Property 1: (2, 1)(6, 5)(7, 3), (2, 1)(6, 5)(7, 4), (2, 1)(6, 5)(4, 3), (2, 1)(4, 3)(6, 5), (2, 1)(4, 3)(7, 6), and (2, 1)(4, 3)(7, 5). The complete set of NJ trees corresponding to Type-3 cones is provided in Figure B3.

**Fig. 3:**
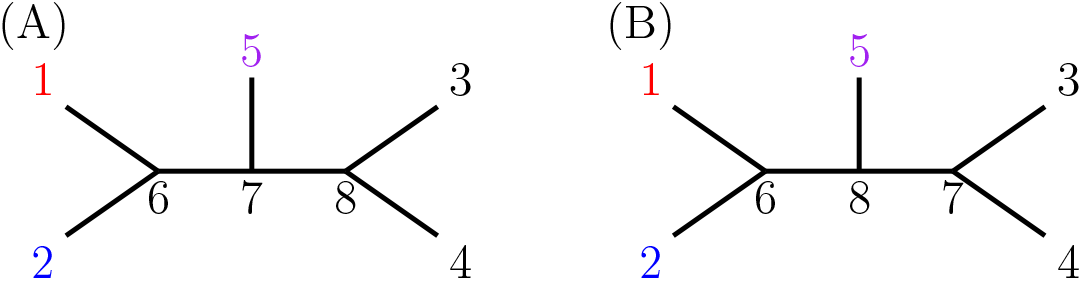
Example NJ tree corresponding to a Type-3 NJ cone violating Property. 1. The two trees shown are the same NJ tree but differ only in their internal node labels, resulting from different cherry-picking orders within the same equivalence class. **(A)** Labeled tree topology, with internal node labeled, from the cherry-picking orders: (2, 1)(6, 5)(7, 3), (2, 1)(6, 5)(7, 4), and (2, 1)(6, 5)(4, 3). **(B)** Labeled tree topology, with internal node labeled, from the cherry-picking orders: (2, 1)(4, 3)(6, 5), (2, 1)(4, 3)(7, 6), and (2, 1)(4, 3)(7, 5).

#### 3.1.2 NJ cones satisfying Property 3 (intermediacy of path length)

Property 3 applies solely to the labeled tree topology of the NJ tree. Since cherry-picking orders determine the labeled tree topology of the final NJ tree, an input dissimilarity vector satisfies this property if and only if it is contained within an NJ cone whose corresponding labeled tree topology satisfies Property 3. Given that all dissimilarity vectors within an NJ cone result in the same labeled tree topology, distinguishing reflection symmetry, either all vectors in the cone satisfy Property 3 or none do. There are sixteen NJ cones whose labeled tree topologies satisfy Property 3. The complete set of trees corresponding to the NJ cones that do not satisfy Property 3 is presented in Figure B4.

#### 3.1.3 Cones satisfying Property 2 (intermediacy of distances)

An NJ cone can contain both dissimilarity vectors that satisfy Property 2 (Section 2.2.2) and those that do not. Therefore, an NJ cone cannot be strictly categorized as either fully satisfying or not satisfying Property 2. Instead, we identify the subset of each NJ cone that satisfies Property 2 and compute its associated volume. To identify the subset of an NJ cone composed exclusively of dissimilarity vectors satisfying Property 2, we must map each input dissimilarity vector **d**^(5)^ to its corresponding tree metric ***δ***, as Property 2 applies to the tree metric of the resulting NJ tree. In this section, we show that there exists a linear transformation from an input dissimilarity vector to its corresponding tree metric and identify cones associated with Property 2.

We first show that, for *N* = 5, the dissimilarity vector **d**^(3)^, obtained after the second cherry-picking step, equals the vector of the minimum path lengths between the two remaining internal nodes and the remaining leaf node. Recall that every cherry-picking order is equivalent to one where the first two cherries consist of the original taxa {1, …, 5} (Section 2.1.5). Thus, we can assume the first cherry corresponds to the leaves on the left side of the final NJ tree, and the second cherry to the right. Under this assumption, the remaining nodes to be clustered are the two internal nodes {6, 7} —formed after selecting the first two cherries—and the remaining taxon from the original set.

##### Lemma 1

Given *N* = 5, let **d**^(3)^ be the dissimilarity vector obtained after the second cherry-picking step. Denote the unpaired original taxon as *e*, and the two newly created nodes as *u* and *v*, respectively. The NJ tree restricted to these three nodes is shown in Figure 4. Then, the following holds:

**Fig. 4:**
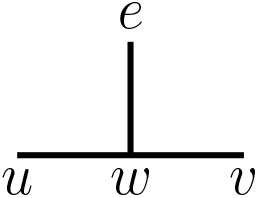
NJ tree restricted to the final three nodes for. *N* = 5. Without loss of generality, for *N* = 5, the first two cherries are assumed to be on opposite sides of the NJ tree. The figure shows the remaining three nodes to be clustered after the first two cherry-picking steps.

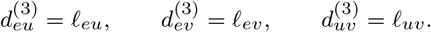

##### Proof.

With three taxa remaining, all the rows of *A*^(3)^ are identical (Section 2.1.5), and thus, any pair from the remaining cherries can be selected in the third cherry-picking step. Without loss of generality, suppose the cherry chosen at the third cherry-picking step is (*e, u*). Then by Eqs. 2, 4, and 5, we have

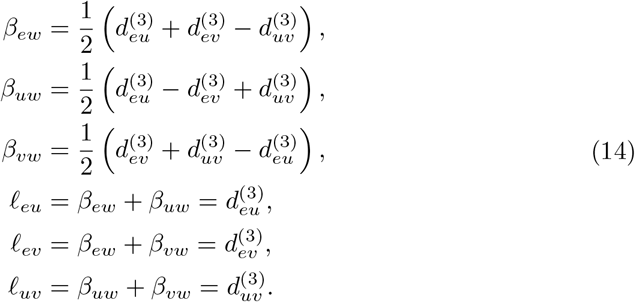

The result also holds when the cherry chosen at the third iteration is either (*e, v*) or (*u, v*); the same proof applies by permuting the node labels {*e, u, v*} accordingly.

We next construct a matrix that maps an input dissimilarity vector to its corresponding tree metric.

##### Definition 8 (Assignment map)

Let *σ* ∈ *S*_5_ be a permutation, where *S*_5_ is the symmetric group on five elements. Consider the unrooted binary tree structure in Figure 5A, where the set of leaves {*a, b, c, d, e*} represents the taxa, and {1, 2, 3, 4, 5} is the index set. Define the *initial assignment ψ*_0_ : {*a, b, c, d, e*} → [5], where each taxon is mapped to its corresponding index as follows:

**Fig. 5:**
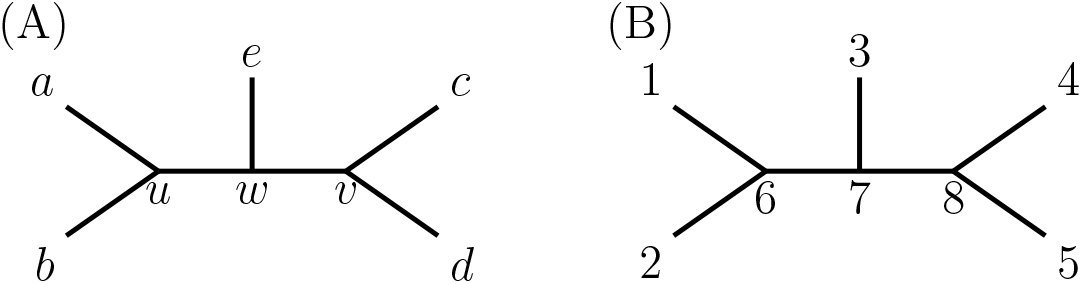
Node labels for an unrooted binary tree for five leaves and their initial assignment. **(A)** The NJ tree for *N* = 5 taxa is shown with leaf labels {*a, b, c, d, e*} and internal nodes {*u, w, v*}. The leaves {*a, b, c, d, e*} correspond to the taxa indexed by {1, 2, 3, 4, 5}. The tree topology remains fixed, while the indices corresponding to each leaf label are permuted according to the *σ*-assignments. **(B)** The leaf labels {*a, b, c, d, e*} are mapped to the taxa indices {1, 2, 3, 4, 5} via the initial assignment *ψ*_0_ (Definition 8). The internal nodes *u, v*, and *w* are labeled as 6, 7, and 8, respectively, according to the cherry-picking order (2, 1)(4, 3)(6, 5).

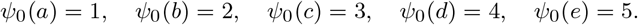

An *σ*-*assignment* is defined as a permutation-induced reassignment of taxa on the tree leaves via *σ* : [5] → [5]. The function *ψ*_*σ*_ = *σ* ∘ *ψ*_0_ describes this reassignment according to *σ*, such that for each *i* ∈ {*a, b, c, d, e*}, *ψ*_*σ*_ (*i*) = *σ*(*ψ*_0_(*i*)).

##### Lemma 2

For a given leaf pair (*ϵ, ζ*) (*ϵ > ζ*) in the labeled tree topology under the initial assignment (Figure 5B), we demonstrate that there exists a column vector ν^(*ϵζ*)^ ∈ ℝ^10×1^ such that the tree metric *δ*_*ϵζ*_is expressed as:

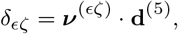

i.e., the shortest path length between the leaves (*ϵ, ζ*) is a linear combination of the entries of the initial dissimilarity vector.

##### *Proof*.

By Definition 8, *ψ*_0_ defines the mapping from the leaves of the NJ tree to the taxa indices. The internal nodes *u, v*, and *w* are assigned to the indices 6, 7, and 8, respectively, under the assumption that the first cherry is (2, 1), the second cherry is 16 (4, 3), and the third cherry is (6, 5). This assumption holds because each equivalence class of cherry-picking orders contains at least one order where the first and second cherries consist solely of the original taxa (Section 2.1.5). Additionally, since all rows of **A**^(3)^ are identical, any remaining pair of nodes can be selected arbitrarily as the third cherry. With this cherry-picking order, the labeled tree topology corresponding to the initial assignment is shown in Figure 5B.

We classify the leaf pairs of the labeled tree topology under the initial assignment into four types based on the structure of the edges along the minimum path between each pair. Type 1 consists of pairs (3, 1), (4, 1), (4, 2), and (3, 2), where each minimum path traverses one edge from the first cherry-picking step, one from the second, and the edges *e*_86_ and *e*_87_. Type 2 includes pairs (5, 3) and (5, 4), where the minimum paths involves edges *e*_85_, *e*_87_, and one edge from the second cherry-picking step. Type 3 includes pairs (5, 1) and (5, 2), whose minimum paths traverse edges *e*_85_, *e*_86_, and one edge from the first cherry-picking step. Finally, Type 4 consists of pairs (2, 1) and (4, 3), where the minimum path length between each pair equals their dissimilarity at the start of the NJ algorithm.

The classification of leaf pairs serves the purpose of expressing the minimum path length, equivalent to the tree metric between leaves, as a linear combination of the entries in the input dissimilarity vector for the five taxa. Two leaf pairs are classified under the same type if their respective linear combinations are related by a permutation of the indices. We claim that if the coefficient vector ***ν***^(*ϵζ*)^ is known for one leaf pair of a given type, the coefficient vector for any other leaf pair of the same type can be computed via a permutation matrix. This follows from the reasoning below.

Let (*η, θ*) (*η > θ*) be another leaf pair of the labeled tree topology with the initial assignment, such that (*η, θ*) belongs to the same type as (*ϵ, ζ*). Define a permutation *σ*′ such that *σ*′(*ϵ*) = *η, σ*′(*ζ*) = *θ, σ*′(*η*) = *ϵ, σ*′(*θ*) = *ζ*, and *σ*′(*χ*) = *χ* for all *χ* ∈ [5] with *χ*≠ *η, ζ, θ, ϵ*. Let *k*_*ij*_ denote the position of the pair (*i, j*) ∈ {(*x, y*) | *x, y* ∈ [5], *x > y*} in the lexicographical ordering. Let 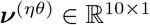 be a column vector where each entry 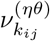equals 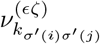 . Let 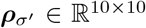 denote the permutation matrix induced by *σ*′, permuting the indices of ***ν***^(*ϵζ*)^. Then,

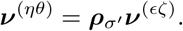

From the classification, leaf pairs of the same type share minimum paths with identical edge compositions, including an equal number of edges from both the first and second cherry-picking steps, as well as edges not involved in either step. Since edges from the same cherry-picking step correspond to the same number of unclustered nodes, their lengths are determined by the same formula, differing only by a permutation of the dissimilarity vector indices. This structural equivalence allows us to compute the tree metric between (*η, θ*) by permuting the known coefficient vector associated with another leaf pair of the same type. Therefore, *δ*_*ηθ*_ is expressed as:

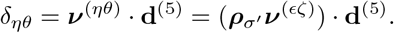

We proceed by constructing ***ν***^(*ϵζ*)^ for a representative pair from each type.

***The first type of pairs consist of*** (3, 1), (4, 1), (4, 2) ***and*** (3, 2).

For the first type, without loss of generality, we express *δ*_31_ in terms of the input dissimilarity vector **d**^(5)^:

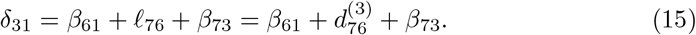

The second step follows from Lemma 1. Using Eqs. 2–5, we compute each term separately as follows:

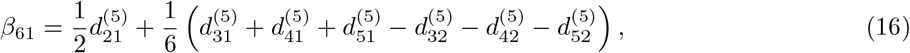

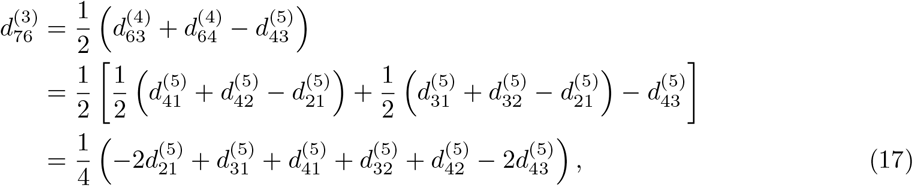

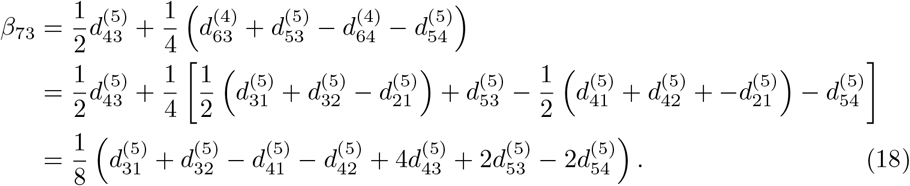

Substituting Eqs. 16–18 into Eq. 15 gives:

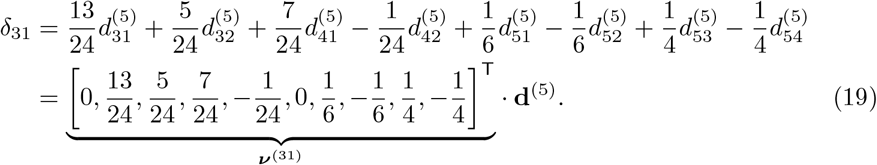

***The second type of pairs consist of*** (5, 3) ***and*** (5, 4).

For the second type, without loss of generality, we express *δ*_53_ in terms of the input dissimilarity vector **d**^(5)^:

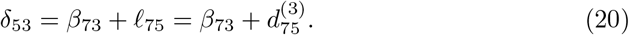

The second step follows from Lemma 1. Using Eqs. 2–5, we compute each term separately as follows:

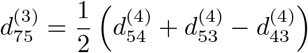

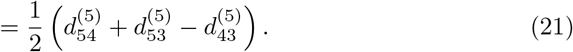

By Eq. 18, we have:

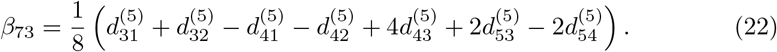

Substituting Eqs. 21 and 22 into Eq. 20 gives:

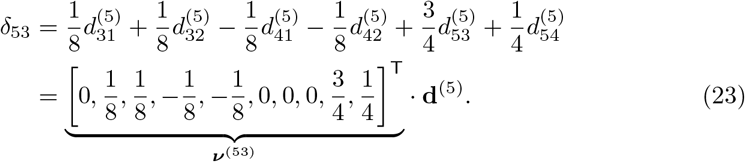

***The third type of pairs consist of*** (5, 1) ***and*** (5, 2).

For the third type, without loss of generality, we express *δ*_51_ in terms of the input dissimilarity vector **d**^(5)^:

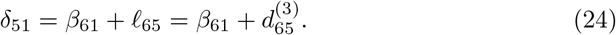

The second step follows from Lemma 1. Using Eqs. 14 and 16, we compute each term separately as follows:

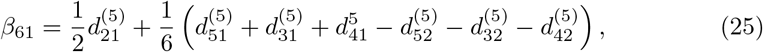

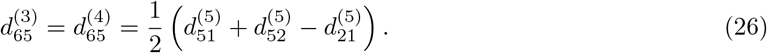

Substituting Eqs. 25 and 26 into Eq. 24 gives:

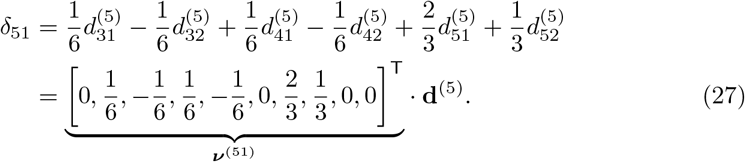

***The fourth type of pairs consist of*** (2, 1) ***and*** (4, 3). From Eq. 5, we have:

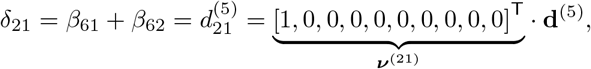

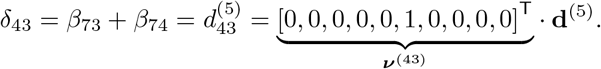

Up to this point, the coefficient vector has been computed for at least one leaf pair in each of the four types. Thus, the tree metric for any leaf pair in the labeled topology, under the initial assignment, can be expressed as a linear combination of the entries in the input dissimilarity vector.

From Eq. 19, the tree metrics for all leaf pairs belonging to the first type are given as follows:

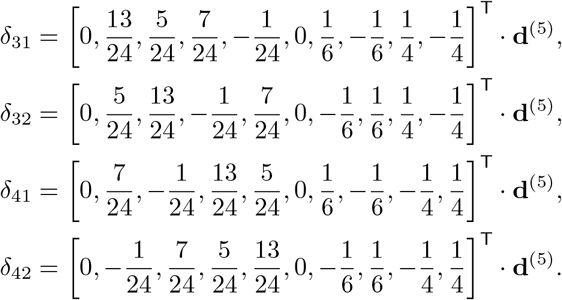

From Eq. 23, the tree metrics for all leaf pairs belonging to the second type are given as follows:

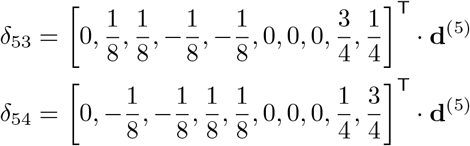

From Eq. 27, the tree metrics for all leaf pairs belonging to the third type are given as follows:

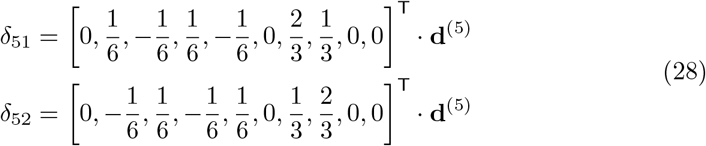

Lemma 2 establishes the existence of a linear transformation from the input dissimilarity vector to the corresponding tree metric, formally stated in the following proposition.

**Proposition 3** For *N* = 5, there exists a linear transformation 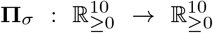, represented by a matrix **Π**_*σ*_, that maps the dissimilarity vector **d**^(5)^ to the tree metric *δ*:

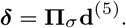

##### *Proof*.

We first construct the matrix **Π**_*id*_ corresponding to the initial assignment *ψ*_0_, where *id* represents the identity permutation. By Eqs. 19, 23, 27–28, the matrix **Π**_*id*_ is defined such that ***δ*** = **Π**_*id*_**d**^(5)^, as follows:

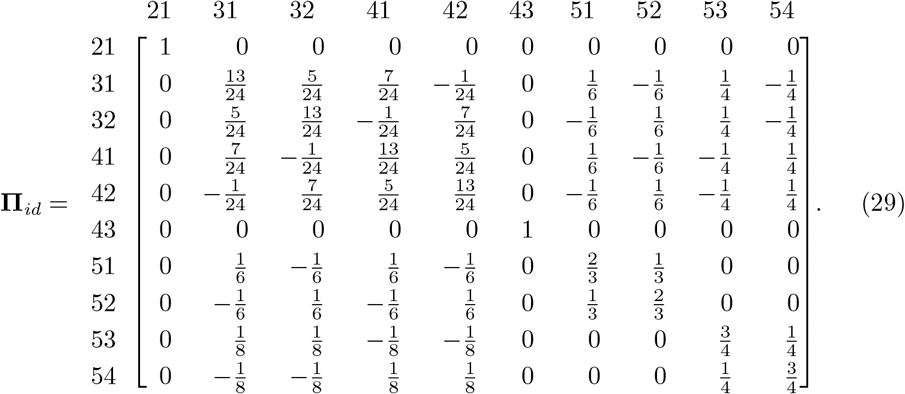

The matrix **Π**_*σ*_ for any *σ*-assignment is constructed by permuting the row and column indices of **Π**_*id*_ according to *σ*. Let *k*_*ij*_ denote the position of the pair (*i, j*) ∈ {(*x, y*) | *x, y* ∈ [5], *x > y*} in the lexicographical ordering. Then the row or column indexed by the pair *k*_*ij*_ is mapped to the row or column indexed by *k*_*σ*(*i*)*σ*(*j*)_.

Formally, let ***ρ***_*σ*_ be the 10 × 10 permutation matrix induced by *σ* acting on the row indices of **Π**_*id*_. The rows of **Π**_*id*_ are permuted by left-multiplication with ***ρ***_*σ*_, while the columns are permuted by right-multiplication with 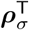. Then, **Π**_*σ*_ is defined as:

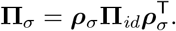

##### Definition 9 (Linear map from dissimilarity vectors to tree metrics for five taxa)

Let **Π**_*id*_ denote the linear map based on the initial assignment, as defined in Eq. 29. The *linear map from dissimilarity vectors to tree metrics for five taxa* **Π**_*σ*_, associated with the *σ*-assignment, is defined as:

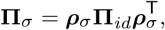

where ρ_*σ*_ is the permutation matrix induced by *σ*. The tree metric *δ*, corresponding to the NJ tree under the *σ*-assignment, is given by:

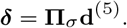

Recall that Property 2 holds if and only if the distance between the admixed taxon and either source taxon is less than the distance between the two source taxa in the final NJ tree. With the tree metric ***δ*** defined as a linear transformation from the input dissimilarity space to the tree metric space (Definition 9), Property 2 is represented by the following matrix:

##### Definition 10 (Tree metric entry comparison matrix)

Define **U** as a linear map from ℝ^10^ to ℝ^2^ given by:

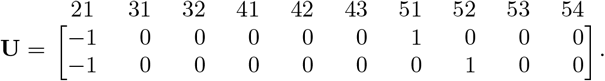

For *N* = 5, the tree metric *δ* satisfies Property 2 if and only if

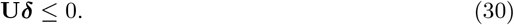

Thus, **U** is referred to as the *tree metric entry comparison matrix*.

By Proposition 3, a tree metric can be expressed as the product of **Π**_*σ*_ and the corresponding input dissimilarity vector. Thus, by Eq. 30, an input dissimilarity vector satisfies Property 2 if and only if the following holds:

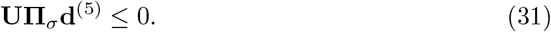

The system of inequalities in Eq. 31 defines a cone bounded by two half-spaces, where each row of the matrix **UΠ**_*σ*_ corresponds to a half-space. To identify the set of dissimilarity vectors in an NJ cone that satisfy Property 2, we intersect the NJ cone with the cone defined by the half-spaces of **UΠ**_*σ*_. For each NJ cone, we augment the half-spaces of **UΠ**_*σ*_ with those defined by the NJ cone matrix in Definition 6. The resulting cone, bounded by this augmented set of half-spaces, is defined as follows:

##### Definition 11 (Property 2 cone)

The *Property 2 cone matrix* 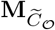 is a 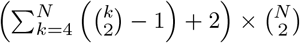 matrix defined as:

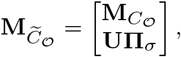

where 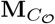 is the NJ cone matrix associated with *C*_𝒪_, **Π**_*σ*_ is the permutation matrix from Proposition 3, and **U** is the comparison matrix enforcing Property 2. For each NJ cone *C*_𝒪_, the *Property 2 cone*, denoted 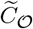, is the subset of *C*_𝒪_ consisting of all input dissimilarity vectors whose corresponding tree metrics satisfy Property 2. It is defined as:

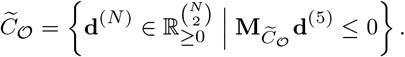

The points within the cones defined by the augmented matrices represent dissimilarity vectors that produce tree metrics satisfying Property 2. Each NJ cone corresponds to a distinct labeled tree topology (not equivalent under reflection about the central node), uniquely determining the assignment of five taxa to the leaves of the final NJ tree. The assignment defines the *σ*-assignment (Definition 8), which allows for the construction of the matrix **Π**_*σ*_. Using the matrices 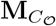 and **U**, we then construct the matrix 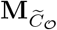 (Definition 11). A dissimilarity vector **d**^(5)^ gives rise to a tree metric satisfying Property 2 if and only if there exists a cherry-picking order whose associated Property 2 cone contains the dissimilarity vector.

### 3.2 Embedding of dissimilarity vectors with admixture

The dissimilarity vector with admixture for *N* = 5, derived from the dissimilarity matrix (Eq. 13), is:

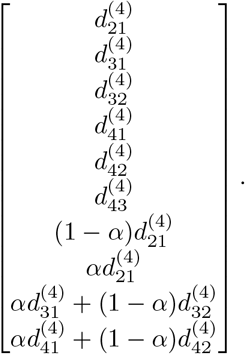

Since the admixed dissimilarity vector contains six independent and four dependent entries, with the latter expressible as linear combinations of the former, there exists an injective linear transformation mapping a dissimilarity vector in ℝ^6^ to an admixed dissimilarity vector in ℝ^10^. Formally, we define:

#### Definition 12 (Embedding map for dissimilarity vector with admixture)

The *embedding map with admixture fraction α*, denoted by *ι*_*α*_ : ℝ^6^ *‘*−→ ℝ^10^, is defined as

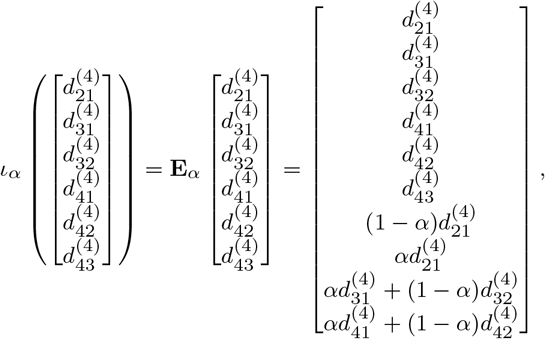

where

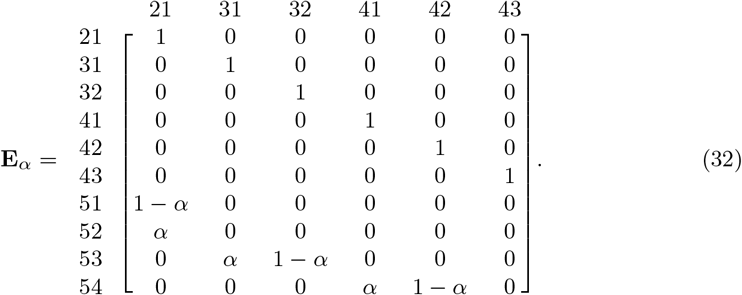

Since the admixed dissimilarity vector involves six independent variables, despite residing in ℝ^10^, we can project NJ cones into ℝ^6^ without loss of information, induced by the inverse of the embedding map *ι*_*α*_. This is formally defined as:

#### Definition 13 (Induced cone)

Let *C* ⊆ ℝ^10^ be a cone, and let **M**_*C*_ ∈ ℝ^*x*×10^ be the matrix whose rows correspond to the inward-pointing normal vectors of the half-spaces defining *C*, where *x* denotes the number of these half-spaces. For a given admixture fraction *α*, define the projection map *π*_*α*_ : ℝ^10^ → ℝ^6^ by the matrix **E**_*α*_ ∈ ℝ^10×6^ (Eq. 32). The image of *C* under *π*_*α*_, denoted *π*_*α*_(*C*) ⊆ ℝ^6^, is the cone whose bounding half-spaces are determined by the rows of the matrix **M**_*C*_ **E**_*α*_. We define *π*_*α*_(*C*) as the *induced cone* by **E**_*α*_.

### 3.3 Computation of cone volumes

In Section 3.1, we identified the cones associated with the three properties for *N* = 5, where every dissimilarity vector in each cone satisfies the corresponding property. Since the dissimilarity vector with admixture has six independent variables, the induced cones reside in ℝ^6^ (Section 3.2). To compute the probability that a specific property holds, we restrict the sample space of dissimilarity vectors to the hypercube [0, 1]^6^, normalizing the maximum dissimilarity between any pair of taxa to 1. This normalization can be achieved by dividing all entries of the dissimilarity vector by its maximum value. Such rescaling is linear and does not affect the outcome of the NJ algorithm. The intersection of each induced cone with this sample space forms a bounded polytope. The probability of the property being satisfied is given by the sum of the volumes of the polytopes formed by the cones satisfying the property, relative to the total volume of the sample space [0, 1]^6^, which is 1.

We constructed the matrices 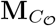 (Definition 6) for all NJ cones *C*_𝒪_ ⊆ ℝ^10^ by generating the matrices **A**^(5)^, **A**^(4)^, and **R** ^(5)^ for each cherry-picking order using the functions `getPairs`, `makeAMatrix`, and `makeRMatrix` from our `NeighborJoining` module implemented in `Macaulay2` (see “Data and Code Availability”). Similarly, we constructed 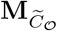 for all Property 2 cones 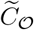 (Definition 11) by generating the matrices **U** and **Π**_*σ*_, then appending them to 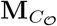 . For a given admixture fraction *α*, we subsequently computed the induced cones (Definition 13), 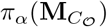 and 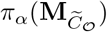 in ℝ using the projection map *π*_*α*_.

We evaluated the volumes of the intersections between the induced cones and the sample space using two approaches. First, we computed the volumes directly using `Macaulay2`. Second, we estimated the volumes via Monte Carlo integration by randomly sampling dissimilarity vectors within a bounded region and calculating the proportion that lies inside the polytope.

#### 3.3.1 Direct volume computation

`Macaulay2` computes the volume of a polytope by first triangulating the polytope into simplices. The triangulation is computed in `Macaulay2` using the software `TOPCOM` [18, 19]. Because there is a closed formula to compute the volume of a simplex that only depends on the vertex matrix of the simplex, it is then straightforward to compute the volume of the polytope as the sum of the volumes of all of these simplices.

Denote the volume of the intersection [0, 1]^6^ ∩ *π*_*α*_(*C*) as 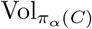, where *C* represents either an NJ cone *C*_𝒪_ or a Property 2 cone 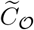. We employed the `Polyhedra` module from `Macaulay2` to compute the exact volumes of these intersections. The function `coneFromHData` was used to transform each **M**_*C*_ into its corresponding cone, which was then mapped to the induced cone *π*_*α*_(*C*). To define the bounding regions, we used `hypercube` to construct the hypercube [−1, 1]^6^ and `posOrthant` to define the positive orthant [0, ∞)^6^. The intersection of these polyhedral components was then computed using `intersect`, and the volume 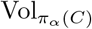 was obtained using the `volume` function. This process provided the exact volumes of [0, 1]^6^ ∩ *π*_*α*_(*C*) for both NJ and Property 2 cones.

Let Ω_all_ represent the set of all equivalence classes of cherry-picking orders, and let Ω_1_ and Ω_3_ denote the sets of equivalence classes of cherry-picking orders satisfying Property 1 and Property 3, respectively (Table 2). The probabilities that a random dissimilarity vector results in an NJ tree satisfying Property 1, Property 2, or Property 3, respectively, are given by:

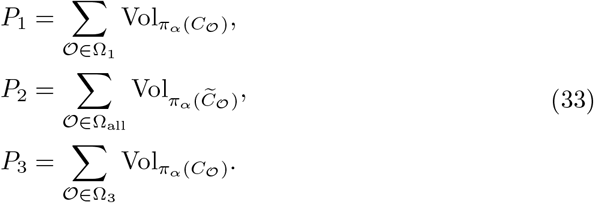

#### 3.3.2 Monte Carlo integration

For each *α* ∈ {0.01, 0.02, …, 0.99}, we generated *N*_sample_ = 100, 000 random vectors **d**^(4)^ ∈ [0, 1]^6^, totalling 99, 00, 000 samples, using the `random` function from `Macaulay2`. Each random vector **d**^(4)^ corresponds to a dissimilarity vector with admixture, **d**^(5)^ = **E**_*α*_**d**^(4)^ ∈ ℝ^10^. For each cone *C* ⊆ ℝ^10^, representing either an NJ cone *C*_𝒪_ or a Property 2 cone 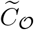, we evaluated whether **d**^(4)^ lies within the induced cone *π*_*α*_(*C*) ⊆ ℝ^6^ by checking the following inequality:

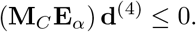

For a given admixture fraction *α*, the probabilities that an NJ tree inferred from **d**^(5)^ = **E**_*α*_**d**^(4)^ satisfies Property 1, Property 2, or Property 3, respectively, are computed as:

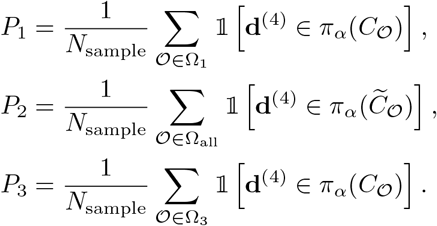

#### 3.3.3 Standard NJ simulation

For comparison with the standard approach [14], we directly applied the NJ algorithm to each randomly generated admixed dissimilarity vector, **d**^(5)^ = **E**_*α*_**d**^(4)^, to infer the corresponding NJ tree using our `NeighborJoining` module in `Macaulay2`. The `runNeighborJoiningClassic` function tracks the cherry-picking order throughout iterations, enabling identification of the corresponding NJ cone. To evaluate Property 1, we assessed whether a cherry containing the two source taxa (1, 2) was selected in the first iteration, as those are the only cones violating Property 1 (Section 3.1.1; Type 3). Properties 2 and 3 were evaluated directly on the inferred NJ trees by first traversing the NJ tree with the `dfs` function to compute the number of edges and shortest path lengths between pairs of source and admixed taxa. These metrics were then compared to test adherence to the respective properties.

## 4 Results

We present the probability that a random dissimilarity vector for *N* = 5 with admixture satisfies the three properties defined in Section 2.2. For each admixture fraction *α* ∈ {0.01, 0.02, …, 0.99}, we computed 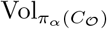 and 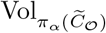, and averaged these values across all *α*. The probabilities of violating the properties were evaluated using three methods: direct volume computation, Monte Carlo integration, and NJ algorithm-based simulations. All methods produced consistent results.

Further analyses revealed the dependency of 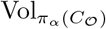 and 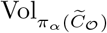 on *α* and its impact on the probabilities of satisfying the defined properties. We provide theoretical insights into why 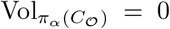 for certain cherry-picking orders and discuss its effect on the probabilities of satisfying Properties 1 and 3. Additionally, we proved the volume equivalence between specific *π*_*α*_(*C*_𝒪_) and their corresponding 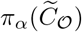, identifying the induced NJ cones that only contain dissimilarity vectors satisfying Property 2.

### 4.1 Mean 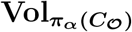 and 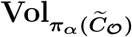 over *α*

Table 2 presents the mean volumes 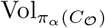 and 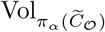 for each 𝒪∈ Ω_all_, averaged across all values of *α* ∈ {0.01, 0.02, …, 0.99}, computed using the direct computation method (Section 3.3.1) For each 𝒪∈ Ω_all_, the induced NJ cones corresponding to the cherry-picking orders indexed 1–10 have volume zero. The proof of this result for all *α* ∈ (0, 1) is provided in Section 4.3. Since 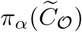 is a subset of *π*_*α*_(*C*_𝒪_), any 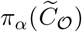 corresponding to a *π*_*α*_(*C*_𝒪_) with volume zero also has a volume zero.

The largest mean 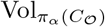 occurs for 𝒪 = (43)(52) and (43)(51) with 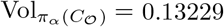. Notably, the mean volumes of the induced Property 2 cones corresponding to these two NJ cones are identical to those of the induced NJ cones themselves, indicating that every dissimilarity vector within these induced NJ cones results in tree metrics that satisfy Property 2. The proof for this result for all *α* ∈ (0, 1) is provided in Section 4.4.

Table 2 groups the thirty labeled tree topologies into six categories, each representing a distinct assignment of two *S* taxa, one *A* taxon, and two *O* taxa to the leaves of the NJ tree. Categories differing only by a reflection about the central node are treated as equivalent. This categorization captures the distinct effects of admixture on the NJ tree structure. The category with the largest mean volume—the average 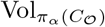 for all 𝒪 within that category—corresponds to cherry-picking orders of the form (*SA*)(*OO*).

### 4.2 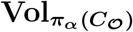 and 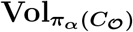 as functions of *α*

Figure 6 presents the volumes of induced NJ cones 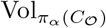, constrained to the sample space [0, 1]^6^, as functions of the admixture fraction *α*, where *α* ∈ {0.01, 0.02, …, 0.99} . The volumes reported are from the direct computation method. *α* ranges from near-boundary values (*α* = 0.01 and *α* = 0.99, where one source population dominates) to the midpoint *α* = 0.5, representing equal contribution from both source populations. Each subplot groups distinct cones based on their mean volume over all *α*-values, as indicated by the NJ cone indices in Table 2. The curves show distinct behaviors depending on the structure of 𝒪, despite the cones in each subplot having identical mean volumes averaged across all admixture fractions. Within each subplot, the NJ cones exhibit two distinct volume trajectories with respect to *α*, even when four cones share the same mean volume.

**Fig. 6:**
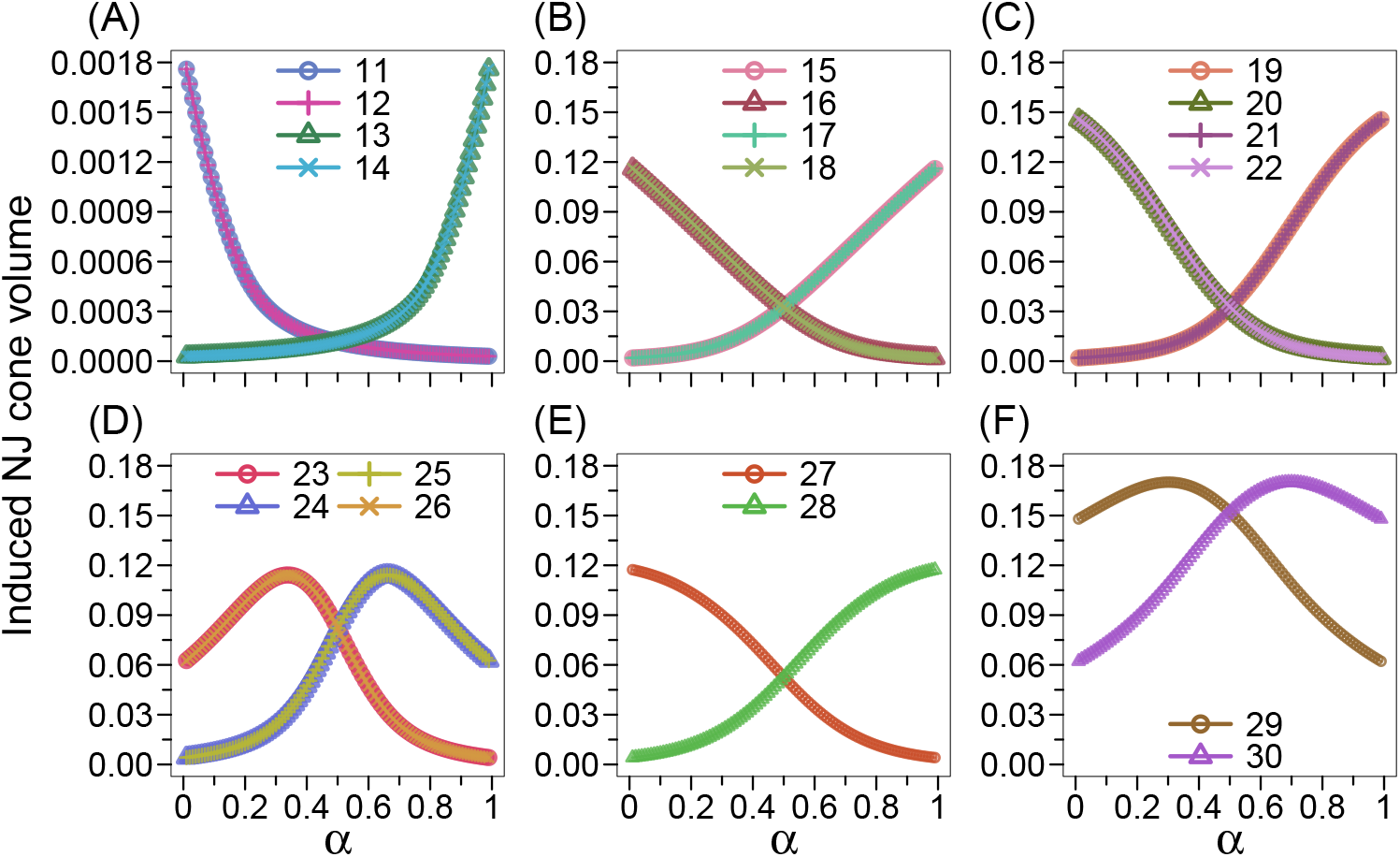
Induced NJ cone volume 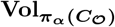 as a function of α. The *x*-axis represents the admixture fraction *α*, where *α* ∈ {0.01, 0.02, …, 0.99}, and the *y*-axis shows the volume of the induced NJ cone constrained to the sample space [0, 1]^6^, i.e., 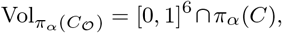, computed using the direct computation method (Section 3.3.1). Each curve corresponds to a distinct NJ cone *C*_𝒪_, indexed as in Table 2 and grouped by mean volume over all *α* values. NJ cones with indices 1–10, for which for 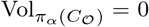 all *α*, are omitted from the figure. NJ cones correspond to: **(A)** (42)(53), (32)(54), (41)(53), and (31)(54); indices 11–14. **(B)** (51)(42), (52)(41), (51)(32), and (52)(31); indices 15–18. **(C)** (42)(51), (41)(52), (32)(51), and (31)(52); indices 19–22. **(D)** (31)(42), (32)(41), (42)(31), and (41)(32); indices 23–26. **(E)** (52)(43) and (51)(43); indices 27, 28. **(F)** (43)(52) and (43)(51); indices 29, 30.

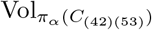 and 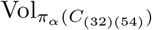 decrease monotonically with *α*, while 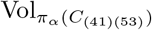 and 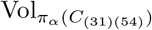 increases over the same range of *α* (Figure 6A). These contrasting monotonic behaviors reflect distinct structural properties of the NJ cones, highlighting the role of admixture fractions in differentiating NJ cone geometries. Cones in the (*O, A*)(*S, O*) category have volumes (maximum: 0.001760227) that are two orders of magnitude smaller than those in Figures 6B–F. The largest volume, 0.1701146, occurs for cones *π*_*α*_(*C*_(43)(52)_) and *π*_*α*_(*C*_(43)(51)_), classified under the (*S, A*)(*O, O*) category (Figures 6F).

Similarly, 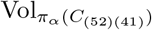 and 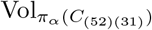 decrease monotonically as *α* increases from 0.01 to 0.99, while 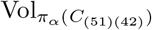 and 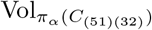 increase monotonically over the same range (Figure 6B). All cherry-picking orders in this subplot are expressed as (*S, A*)(*S, O*), where the first cherry consists of a source taxon and an admixed taxon. For small 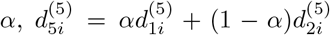 (Eq. 12), which makes 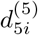 closer to 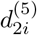, increasing the probability of selecting (5, 2) as the first cherry. Conversely, for large 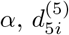 becomes closer to 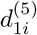 than to 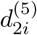, favoring the selection of (5, 1) as the first cherry. This analogous pattern is observed for cones in panels C and E.

Figures 6D and F exhibit a qualitative difference from the other panels, where two volumes increase while two decrease monotonically with *α*. In contrast, Figures 6D and F show more complex, non-monotonic relationships between *α* and the induced NJ cone volumes. In Figure 6D, cones *π*_*α*_(*C*_(31)(42)_) and *π*_*α*_(*C*_(41)(32)_) attain their maximum and minimum volumes at *α* = 0.34 and *α* = 0.99, respectively, while cones *π*_*α*_(*C*_(32)(41)_) and *π*_*α*_(*C*_(42)(31)_) reach their maximum at *α* = 0.66 and minimum at *α* = 0.01. All these cones share the same maximum volume of 0.1142092 and minimum volume of 0.001760227. In Figure 6F, cone *π*_*α*_(*C*_(43)(52)_) reaches its maximum volume at *α* = 0.30 and minimum volume at *α* = 0.99, while cone *π*_*α*_(*C*_(43)(51)_) attains its maximum at *α* = 0.70 and minimum at *α* = 0.01. Both cones share the largest maximum volume of 0.1701146 and the largest minimum volume of 0.06219378 across all cones for any *α*.

Figure 7 shows the induced Property 2 cone volumes 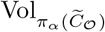 as a function of *α*. The results qualitatively align with Figure 6, except for Figure 7C, though the volumes are consistently smaller for each 𝒪 and *α* due to the induced Property 2 cone being a subset of the corresponding induced NJ cone. Unlike Figure 6C, Figure 7C does not exhibit strict monotonicity with respect to *α*. The volumes of *π*_*α*_(*C*_(42)(51)_) and *π*_*α*_(*C*_(32)(51)_) increase monotonically until *α* = 0.91, reaching a maximum of 0.09632147, after which they decrease monotonically as *α* approaches 0.99. Conversely, the volumes of *π*_*α*_(*C*_(41)(52)_) and *π*_*α*_(*C*_(31)(52)_) follow an opposite trend, increasing monotonically until *α* = 0.09, reaching the same maximum of 0.09632147, and decreasing monotonically for *α* ∈ (0.09, 0.99).

**Fig. 7:**
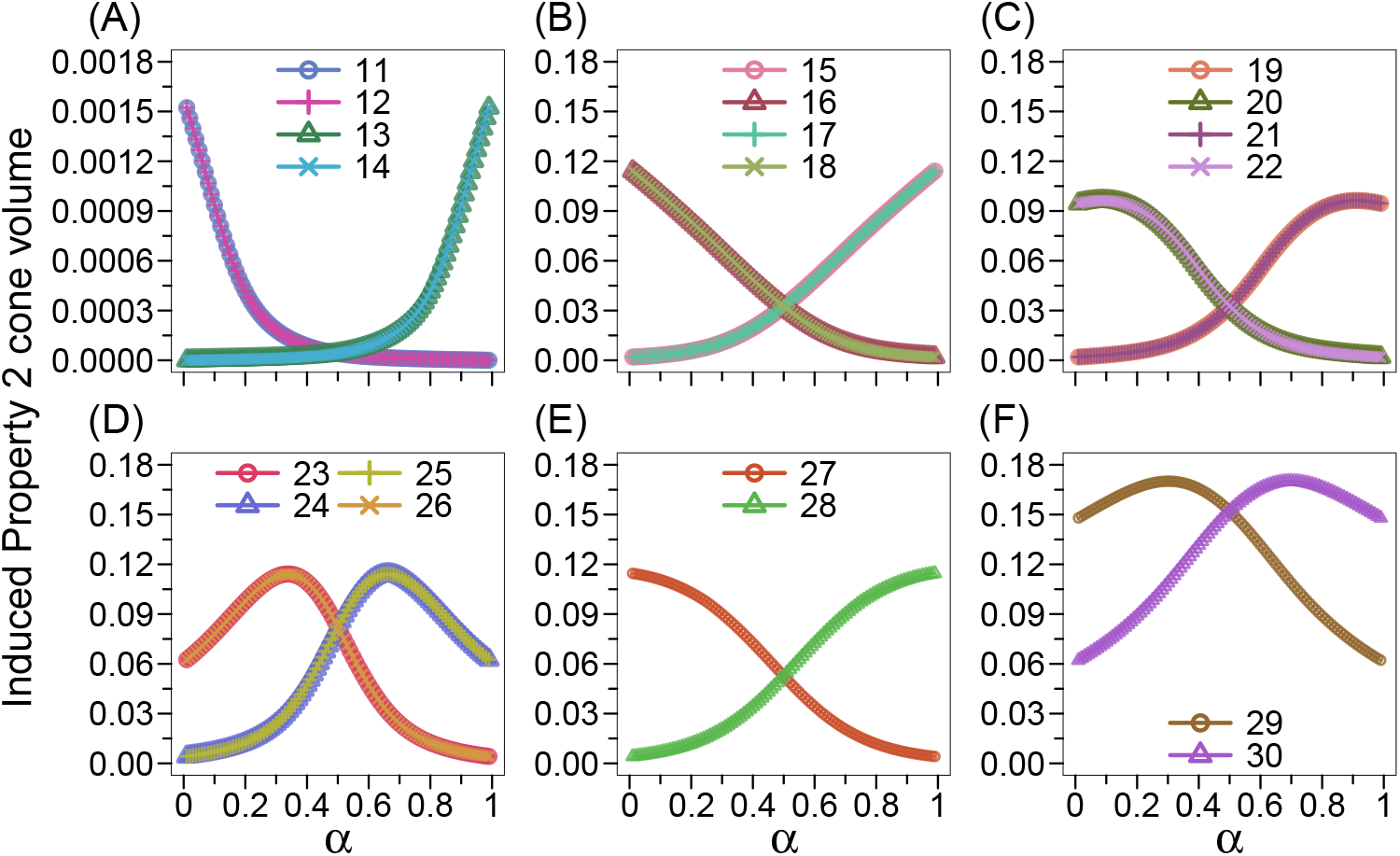
Induced Property 2 cone volume 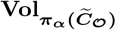 as a function of α. The *x*-axis represents the admixture fraction *α*, where *α* ∈ {0.01, 0.02, …, 0.99}, and the *y*-axis shows the volume of the induced Property 2 cone constrained to the sample space [0, 1]^6^, i.e., 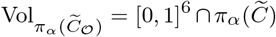, computed using the direct computation method (Section 3.3.1). The cone indices follow those listed in Table 2. NJ cones with indices 1–10, for which 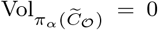 for all *α*, are omitted from the figure. The figure layout mirrors that of Figure 6. Property 2 cones correspond to: **(A)** (42)(53), (32)(54), (41)(53), and (31)(54); indices 11–14. **(B)** (51)(42), (52)(41), (51)(32), and (52)(31); indices 15–18. **(C)** (42)(51), (41)(52), (32)(51), and (31)(52); indices 19–22. **(D)** (31)(42), (32)(41), (42)(31), and (41)(32); indices 23–26. **(E)** (52)(43) and (51)(43); indices 27, 28. **(F)** (43)(52) and (43)(51); indices 29, 30.

Table A1 reports the computed volumes 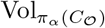 and 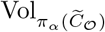 for each 𝒪at specific values of *α* = 0.01, 0.5, and 0.99. These *α*-values were chosen to capture the behavior at the extremities of the parameter space, where *α* = 0.01 and *α* = 0.99 correspond to near-complete contribution from a single source population, and *α* = 0.5 represents an equal admixture scenario.

### 4.3 Cherry-picking orders with 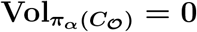

Both the direct volume computation method and the Monte Carlo method support the following proposition, which we prove using two independent methods.

**Proposition 4** For any admixture fraction *α* ∈ (0, 1), 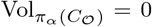 if 𝒪corresponds to one of the following cherry-picking orders:

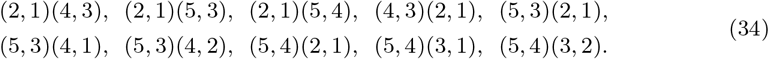

*Proof*. ***Method 1: proof by contradiction***

Let 𝒪be a cherry-picking order from Eq. 34. By applying the Fourier–Motzkin elimination [20–22] to the system of inequalities 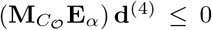, which defines the induced NJ cone *π*_*α*_(*C*_𝒪_), we obtain that there exists *i* ∈ [6] such that 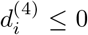 [14]. This contradicts the assumption that all entries of **d**^(4)^ are strictly positive. Thus, no dissimilarity vector **d**^(4)^ belongs to the induced cone *π*_*α*_(*C*_𝒪_), implying its volume is 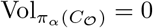.

***Method 2: proof by dimensionality***

The dimensions of the induced NJ cones *π*_*α*_(*C*_𝒪_) were computed in `Macaulay2`. All cones corresponding to the cherry-picking orders in Eq. 34 have dimensions strictly less than 6, indicating that they reside in proper subspaces of ℝ^6^. Since the dimension of each cone is strictly lower than that of the ambient space, their volumes in ℝ^6^ are zero.

#### Corollary 4.1

For any admixture fraction *α* ∈ (0, 1), the probability that an admixed dissimilarity vector **d**^(5)^ violates Property 1 is zero.

*Proof*. Only Type-3 cherry-picking orders violate Property 1 (Section 3.1.1), and Proposition 4 lists all such Type-3 cherry-picking orders, showing that the induced cones corresponding to these orders have zero volume. Thus, for a dissimilarity vector **d**^(5)^ with admixture, no Type-3 cherry-picking orders are possible. Therefore, the probability of **d**^(5)^ violating Property 1 is zero.

#### Corollary 4.2

For any admixture fraction *α* ∈ (0, 1), the probability that an admixed dissimilarity vector **d**^(5)^ violates Property 3 is given by the sum of volumes of the induced NJ cones corresponding to the following cherry-picking orders:

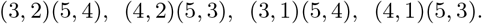

#### *Proof*.

There are 14 cherry-picking orders that violate Property 3 (Section 3.1.2). Of these, 10 correspond to volume-zero induced NJ cones as listed in Proposition 4, while the remaining four have non-zero volumes. Thus, the probability of violating Property 3 is given by the sum of the volumes of these four *π*_*α*_(*C*_𝒪_)’s not included in Proposition 4.

### 4.4 Cherry-picking orders with 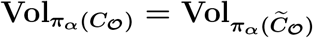

**Proposition 5** For any admixture fraction *α* ∈ (0, 1), no dissimilarity vector **d**^(4)^ within the induced NJ cones *π*_*α*_(*C*_(4,3)(5,1)_) and *π*_*α*_(*C*_(4,3)(5,2)_) (Figure B5) violates Property 2.

*Proof*. We proceed by contradiction, assuming that Property 2 is violated. This implies that either *δ*_21_ *< δ*_51_ or *δ*_21_ *< δ*_52_.

The *σ*-assignment corresponding to the induced NJ cone *π*_*α*_(*C*_(4,3)(5,1)_), and thereby the labeled tree topology in Figure B5A, reorders the taxa from the initial assignment (Figure 5B) as follows:

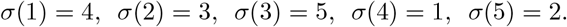

The linear map from dissimilarity vectors to tree metrics for this *σ*-assignment, 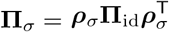 (Definition 9), is then given by:

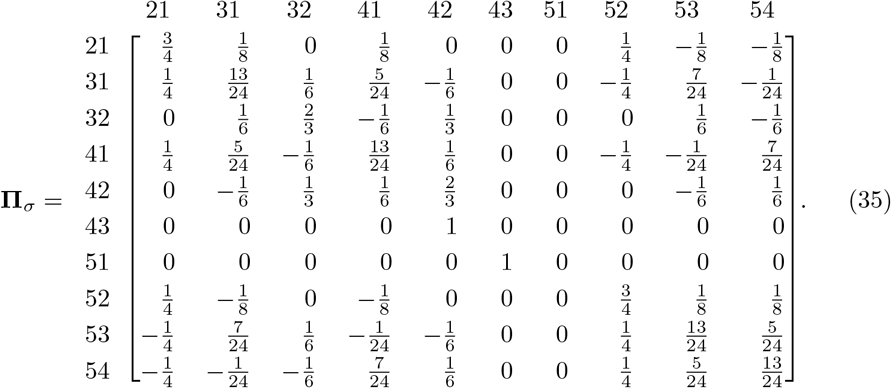

We first consider the case where 𝒪= (4, 3)(5, 1) and *δ*_21_ *< δ*_52_. Then by Eq. 35,

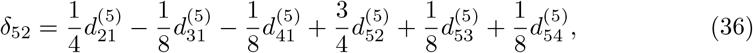

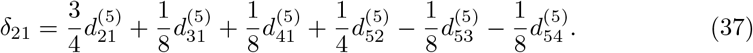

Subtracting Eq. 36 from Eq. 37 results in the following:

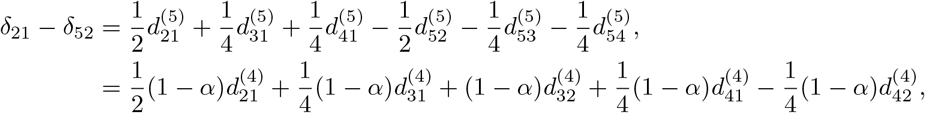

where the last step follows from **d**^(5)^ = **E**_*α*_**d**^(4)^ (Definition 12). Then *δ*_21_ *< δ*_52_ can be written as the following inequality:

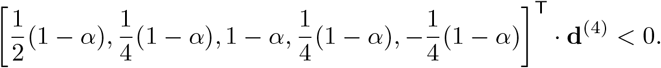

Dividing both sides of the inequality by 1 − *α* gives:

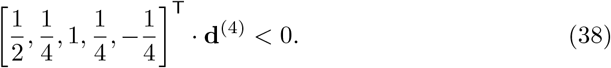

By Definition 13, every dissimilarity vector **d**^(4)^ contained in *π*_*α*_(*C*_(4,3)(5,1)_) satisfies:

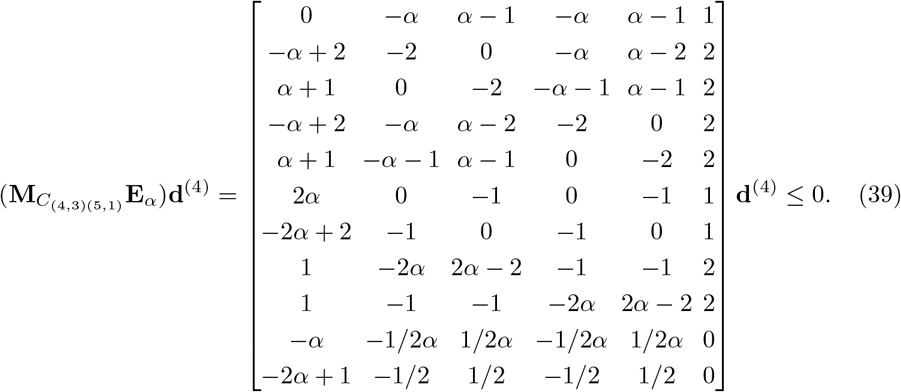

Multiplying Eq. 38 by 2 and adding it to the last row of Eq. 39 yields:

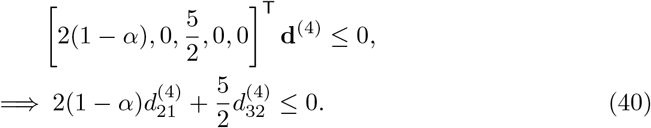

Since 2(1 − *α*) *>* 0 and 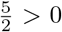, Eq. 40 implies that either 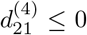 or 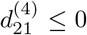, which contradicts the assumption that all entries of **d**^(4)^ are strictly positive.

In the second case,𝒪 = (4, 3)(5, 1) and *δ*_21_ *< δ*_51_. As in the first case, if the dissimilarity vector **d**^(5)^ = **E**_*α*_**d**^(4)^ corresponds to a tree metric where *δ*_21_ *< δ*_51_, the following inequality holds:

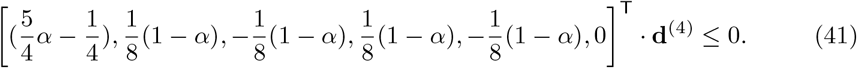

Multiplying Eq. 41 by 8 and adding it to the last row of Eq. 39, scaled by 2(1 − *α*), results in the following:

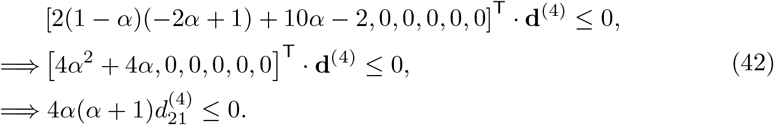

Since 4*α >* 0 and *α* + 1 *>* 0, Eq. 42 implies 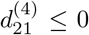, contradicting the assumption cthat all entries of **d**^(4)^ are strictly positive.

Thus, no dissimilarity vector in *π*_*α*_(*C*_(4,3)(5,1)_) violates Property 2. By analogous reasoning, the same conclusion holds for dissimilarity vectors in *π*_*α*_(*C*_(4,3)(5,2)_), which corresponds to the labeled tree topology in Figure B5B.

### 4.5 Probability of satisfying each of the three properties

For an admixed dissimilarity vector with *N* = 5, the probability of satisfying Property 1 is always 1, independent of *α*. Only Type-3 equivalence classes corresponding to the cherry-picking orders (21)(54), (21)(53), and (21)(43) violate Property 1 (Section 3.1.1). Two independent approaches confirmed that the NJ cones corresponding to these equivalence classes of cherry-picking orders have zero volume. The first approach involves direct computation from Table 2, and the second follows from Proposition 4 in Section 4.3.

The probability of satisfying Property 2 equals the total volume of all cones satisfying Property 2 (Eq. 33). Figure 8A shows how *P*_2_ depends on the admixture parameter *α*, computed via direct volume computation. For *α* = 0.5, representing equal contribution from both source populations, the probability reaches its maximum value of 0.9996976. As *α* deviates from 0.5, reflecting increasing asymmetry in the admixture proportions, the probability decreases symmetrically, reaching a minimum of 0.8903937 at the extreme values *α* = 0.01 and 0.99, where one source population contributes almost entirely to the admixed population. The observed symmetry around *α* = 0.5 aligns with the interchangeable roles of the two source populations in the assumed admixture model.

**Fig. 8:**
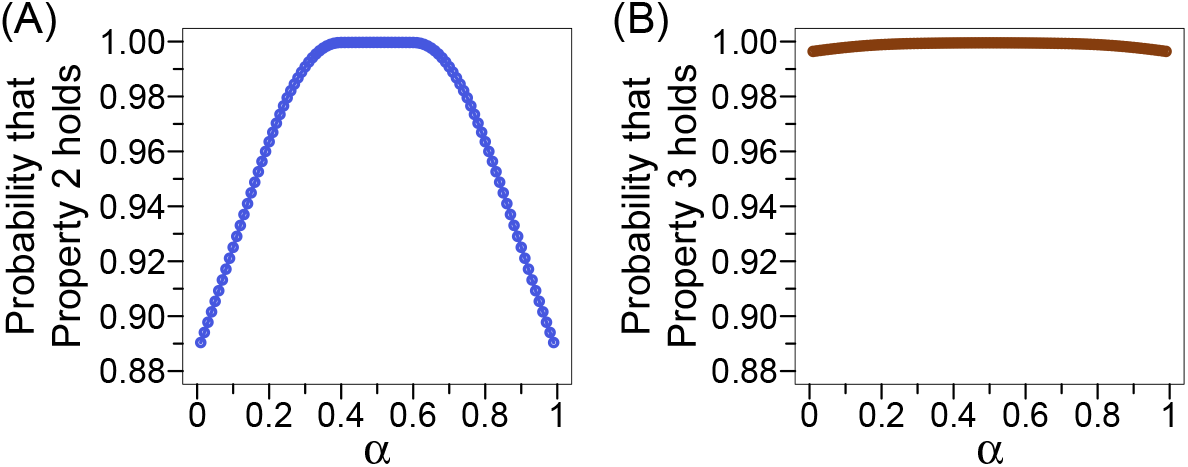
*P*_2_ and *P*_3_ as functions of α. The *x*-axis denotes the admixture fraction *α*, where *α* ∈ {0.01, 0.02, …, 0.99}, and the *y*-axis represents the probability that a given property holds, computed using the direct computation method (Section 3.3.1). For each value of *α*, the probability is obtained by summing the volumes of the induced cones intersecting with the sample space [0, 1]^6^. **(A)** The probability that Property 2 holds, 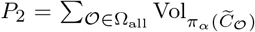, where the summation is taken over all cherry-picking orders. **(B)** The probability that Property 3 holds, 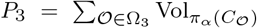, where the summation is restricted to the cones corresponding to cherry-picking orders that satisfy Property 3. Property 1 holds for all admixed dissimilarity vectors **d**^(5)^ = **E**_*α*_**d**^(4)^ across all values of *α*, i.e., *P*_1_ = 1 for all *α* ∈ (0, 1) (Corollary 4.1). Thus, the corresponding plot for Property 1 is omitted from the figure.

Similar qualitative behavior is observed for Property 3 (Figure 8B), whose probability is computed as the total volume of the induced NJ cones satisfying Property 3 (Eq. 33, Figure B4). As with Property 2, the maximum probability occurs at balanced admixture (*α* = 0.5) with a value of 0.999537, which is slightly higher than that of Property 2. However, unlike Property 2, even at extreme admixture proportions (*α* = 0.01 and *α* = 0.99), the probability remains high, at 0.999537. This indicates that Property 3 is robust to skewed admixture, while Property 2 is more sensitive to asymmetry in the contributions from the source populations.

To assess the accuracy of the direct volume computation and Monte Carlo methods, we compared their results with those obtained from the standard NJ simulation [14] in `Macaulay2` (Section 3.3). Table 3 summarizes the probabilities of violating each of the three properties across the methods. The results show close agreement, with differences only at the fifth decimal place.

**Table 3:**
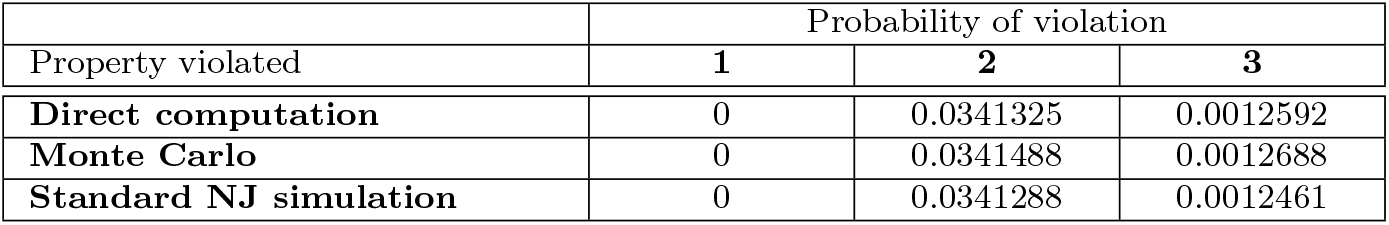
Methods comparison. This table shows the probabilities of violating each of the three properties averaged across all *α* ∈ {0.01, 0.02, …, 0.99}. In the direct computation method (Section 3.3.1), the probability of violation was computed as 1 minus the total volume of the induced cones satisfying each property within the sample space [0, 1]^6^. In the Monte Carlo method (Section 3.3.2), 100,000 dissimilarity vectors **d**^(4)^ were uniformly sampled from [0, 1]^6^ for each of the 99 *α* values. The violation probability was computed as the proportion of vectors falling outside the induced cones, based on the total of 9,900,000 samples. In the standard NJ simulation (Section 3.3.3), the NJ algorithm was directly applied to each dissimilarity vector **d**^(5)^ = **E**_*α*_**d**^(4)^ to infer the corresponding NJ tree. The probability of violation was then calculated as the fraction of NJ trees (out of 9,900,000) that failed to satisfy the properties.

| Property violated | Probability of violation |  |  |
| --- | --- | --- | --- |
|  | 1 | 2 | 3 |
| **Direct computation** | 0 | 0.0341325 | 0.0012592 |
| **Monte Carlo** | 0 | 0.0341488 | 0.0012688 |
| **Standard NJ simulation** | 0 | 0.0341288 | 0.0012461 |

## 5 Discussion

In this study, we have investigated the geometric and probabilistic behavior of the NJ algorithm applied to distance matrices under admixture, focusing on a five-taxon case. By formulating the problem using polyhedral cones and projection maps, we have introduced a geometric framework that partitions the space of dissimilarity vectors based on their clustering, distance, and topological path length properties. This approach has enabled direct computation of the associated probabilities by evaluating the volumes of the induced cones within the bounded dissimilarity space. We have validated our analytical results via Monte Carlo integration and classical NJ simulations, confirming their accuracy. We have shown that while Property 1 is always satisfied, the probabilities of satisfying Properties 2 and 3 depend significantly on the admixture fraction. We have also proven that certain induced NJ cones have zero volume when admixture is present, indicating that these topologies are structurally incompatible with admixture under the NJ framework. Our study contributes to advancing the theoretical understanding of how admixture affects the NJ tree inference. Further, our `Macaulay2` implementation enables efficient analysis of the NJ algorithm under admixture, providing a valuable tool for studying complex evolutionary relationships involving admixed populations.

Although our analysis has focused on the five-taxon case, the geometric and combinatorial structure of NJ cones and their projections to lower-dimensional spaces can be generalized to cases with *N >* 5 taxa. Our framework can also be extended to multi-way linear admixture models beyond two-way admixture, introducing additional constraints on the dissimilarity vectors. These complexities can be addressed by extending the projection map to accommodate higher-order mixtures and defining new classes of induced NJ cones that capture the more complex admixture relationships among taxa. Beyond linear distance measures, further theoretical exploration should focus on formalizing the behavior of the NJ algorithm under non-linear genetic distances, such as *F*_ST_ [23] and other *F* -statistics [24, 25], which would require modifications to the current linear projection map.

The NJ algorithm has been shown to be a greedy heuristic [26] for the balanced minimum evolution (BME) problem [27, 28]. A natural extension of our current geometric approach would be to investigate whether the same properties involving admixture and associated probabilities observed under NJ also hold in BME. By leveraging the geometric methods from this work, we can analyze the BME cones [15, 16, 29] analogous to the NJ cones. Such an analysis would reveal structural similarities and differences between NJ and BME, providing a deeper understanding of how algorithmic choices impact phylogenetic tree construction under complex evolutionary models, including cases of admixture.

Finally, our framework provides a principled approach for studying an admixed taxon as a “rogue taxon” [30, 31]. Kim et al. [14] demonstrated via simulation that the three properties hold more frequently when the distances among *N* − 1 non-admixed taxa are additive, a phenomenon linked to the rogue taxon behavior. Since a metric is additive if and only if it satisfies the four-point condition [2], defined by a set of linear inequalities, our geometric framework is ideally suited to rigorously analyze how the addition of an admixed taxon to an underlying set of *N* − 1 populations with a tree-like evolutionary history affects the topological stability of inferred source trees.

## Data and Code Availability

All code used in this manuscript, including our implementation of the NJ algorithm as the module `NeighborJoining`, is available on Zenodo at 10.5281/zenodo.13363307.

## Acknowledgements

This work was supported by National Science Foundation grant DMS-2001367.

## Appendix A Supplementary Tables

**Table A1:**
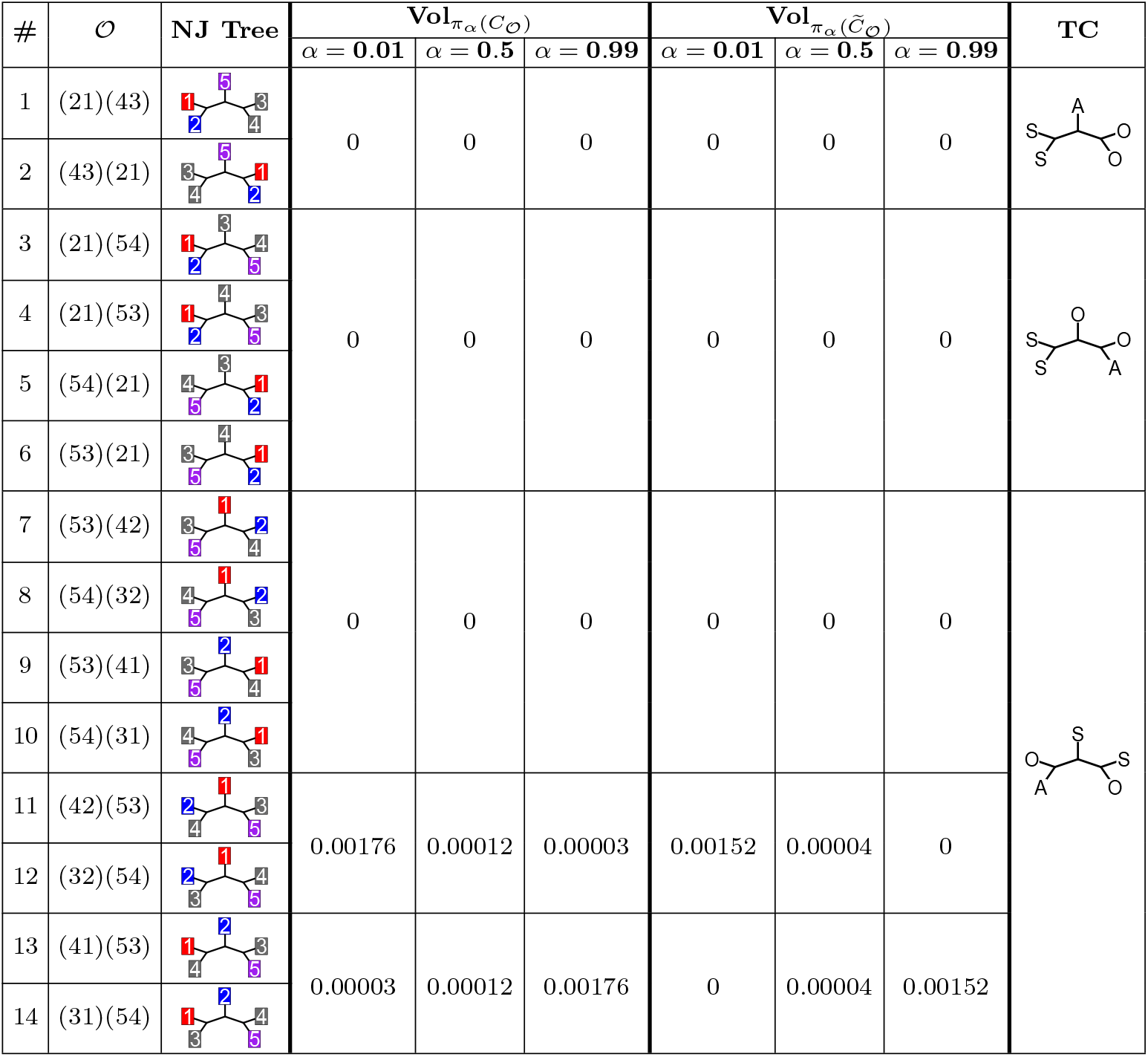
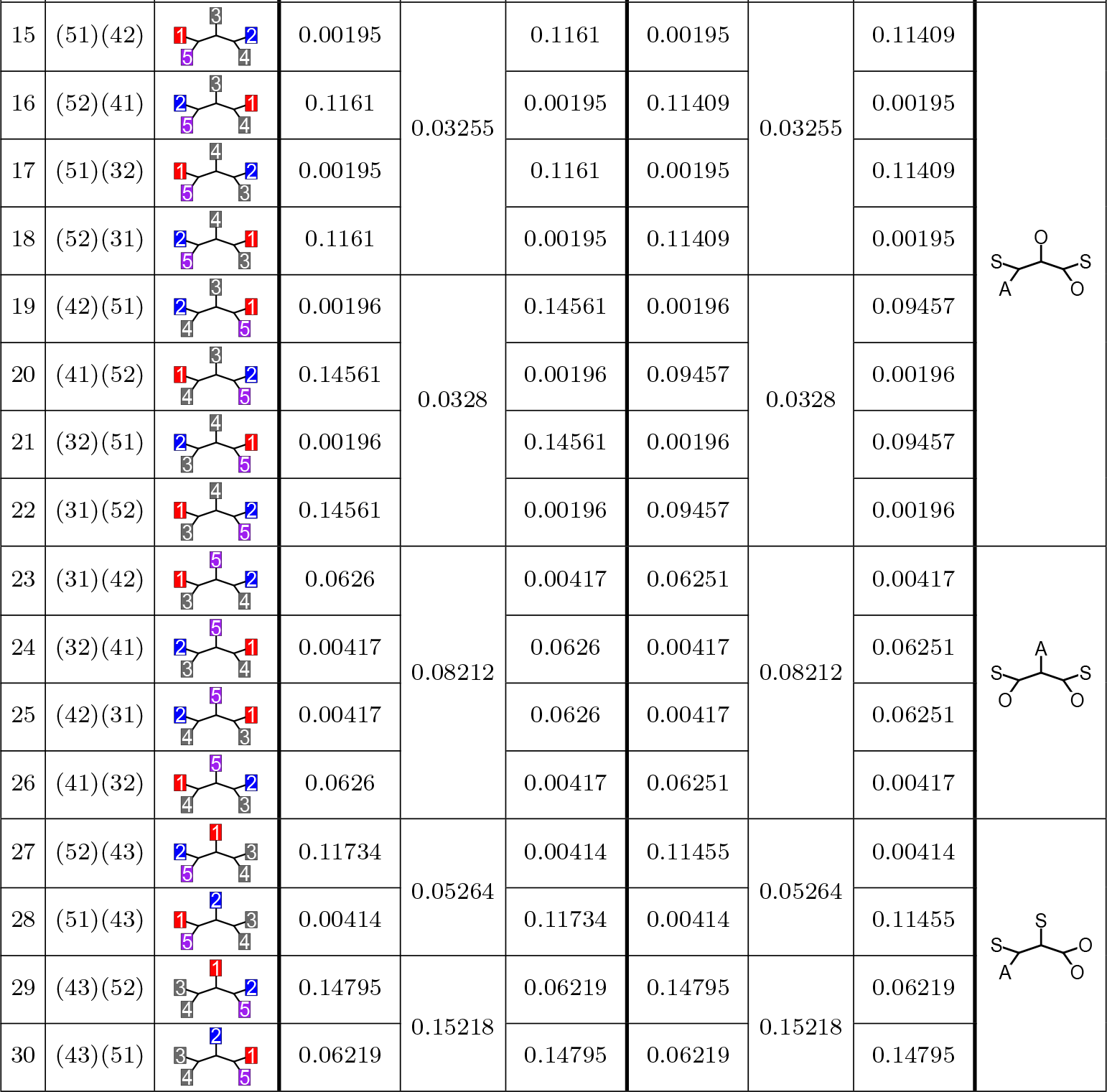
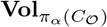 and 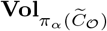 for α = 0.01, 0.5 and 0.99. The table mirrors the structure of Table 2, with mean volumes replaced by those computed for fixed values of *α* = 0.01, 0.5, and 0.99. For each fixed *α* and cherry-picking order 𝒪∈ Ω, the volumes 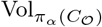 and 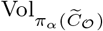 were obtained using the direct computation method (Section 3.3.1). The final column, labeled “TC”, stands for “Tree Category”.

| # | $\mathcal{O}$ | NJ Tree | $\text{Vol}_{\pi_{\alpha}(C_{\mathcal{O}})}$ | | | $\text{Vol}_{\pi_{\alpha}(\bar{C}_{\mathcal{O}})}$ | | | TC | |
| --- | --- | --- | --- | --- | --- | --- | --- | --- | --- | --- |
| | | | $\alpha = 0.01$ | $\alpha = 0.5$ | $\alpha = 0.99$ | $\alpha = 0.01$ | $\alpha = 0.5$ | $\alpha = 0.99$ | | |
| 1 | (21)(43) |  | 0 | 0 | 0 | 0 | 0 | 0 |  |  |
| 2 | (43)(21) |  |  |  |  |  |  |  |  |  |
| 3 | (21)(54) |  | 0 | 0 | 0 | 0 | 0 | 0 |  |  |
| 4 | (21)(53) |  |  |  |  |  |  |  |  |  |
| 5 | (54)(21) |  |  |  |  |  |  |  |  |  |
| 6 | (53)(21) |  |  |  |  |  |  |  |  |  |
| 7 | (53)(42) |  | 0 | 0 | 0 | 0 | 0 | 0 |  |  |
| 8 | (54)(32) |  |  |  |  |  |  |  |  |  |
| 9 | (53)(41) |  |  |  |  |  |  |  |  |  |
| 10 | (54)(31) |  |  |  |  |  |  |  |  |  |
| 11 | (42)(53) |  | 0.00176 | 0.00012 | 0.00003 | 0.00152 | 0.00004 | 0 |  |  |
| 12 | (32)(54) |  |  |  |  |  |  |  |  |  |
| 13 | (41)(53) |  | 0.00003 | 0.00012 | 0.00176 | 0 | 0.00004 | 0.00152 |  |  |
| 14 | (31)(54) |  |  |  |  |  |  |  |  |  |

| # | $\mathcal{O}$ | NJ Tree | $\text{Vol}_{\pi_{\alpha}(C_{\mathcal{O}})}$ | | | $\text{Vol}_{\pi_{\alpha}(\tilde{C}_{\mathcal{O}})}$ | | | TC |
| --- | --- | --- | --- | --- | --- | --- | --- | --- | --- |
| | | | $\alpha = 0.01$ | $\alpha = 0.5$ | $\alpha = 0.99$ | $\alpha = 0.01$ | $\alpha = 0.5$ | $\alpha = 0.99$ | |
| 15 | (51)(42) |  | 0.00195 | 0.03255 | 0.1161 | 0.00195 | 0.03255 | 0.11409 |  |
| 16 | (52)(41) |  | 0.1161 |  | 0.00195 | 0.11409 |  | 0.00195 |  |
| 17 | (51)(32) |  | 0.00195 |  | 0.1161 | 0.00195 |  | 0.11409 |  |
| 18 | (52)(31) |  | 0.1161 |  | 0.00195 | 0.11409 |  | 0.00195 |  |
| 19 | (42)(51) |  | 0.00196 | 0.0328 | 0.14561 | 0.00196 | 0.0328 | 0.09457 |  |
| 20 | (41)(52) |  | 0.14561 |  | 0.00196 | 0.09457 |  | 0.00196 |  |
| 21 | (32)(51) |  | 0.00196 |  | 0.14561 | 0.00196 |  | 0.09457 |  |
| 22 | (31)(52) |  | 0.14561 |  | 0.00196 | 0.09457 |  | 0.00196 |  |
| 23 | (31)(42) |  | 0.0626 | 0.08212 | 0.00417 | 0.06251 | 0.08212 | 0.00417 |  |
| 24 | (32)(41) |  | 0.00417 |  | 0.0626 | 0.00417 |  | 0.06251 |  |
| 25 | (42)(31) |  | 0.00417 |  | 0.0626 | 0.00417 |  | 0.06251 |  |
| 26 | (41)(32) |  | 0.0626 |  | 0.00417 | 0.06251 |  | 0.00417 |  |
| 27 | (52)(43) |  | 0.11734 | 0.05264 | 0.00414 | 0.11455 | 0.05264 | 0.00414 |  |
| 28 | (51)(43) |  | 0.00414 |  | 0.11734 | 0.00414 |  | 0.11455 |  |
| 29 | (43)(52) |  | 0.14795 | 0.15218 | 0.06219 | 0.14795 | 0.15218 | 0.06219 |  |
| 30 | (43)(51) |  | 0.06219 |  | 0.14795 | 0.06219 |  | 0.14795 |  |

## Appendix B Supplementary Figure

**Fig. B1:**
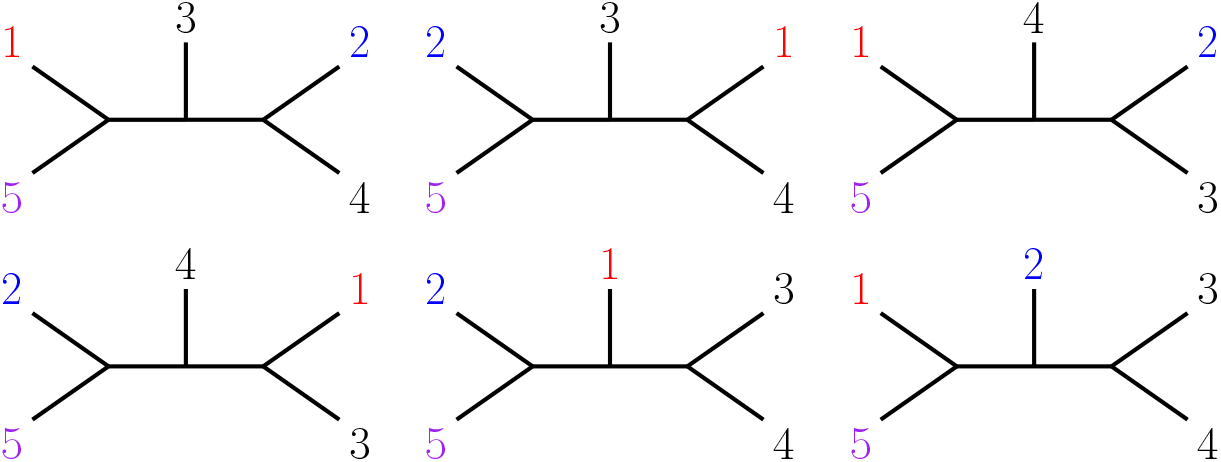
Type 1 labeled tree topologies for Property 1. These six labeled tree topologies correspond to Type-1 NJ cones (Section 3.1.1), where each equivalence class consists entirely of cherry-picking orders that satisfy Property 1. The labeled tree topologies are distinguished based on reflection symmetry with respect to the central taxon.

**Fig. B2:**
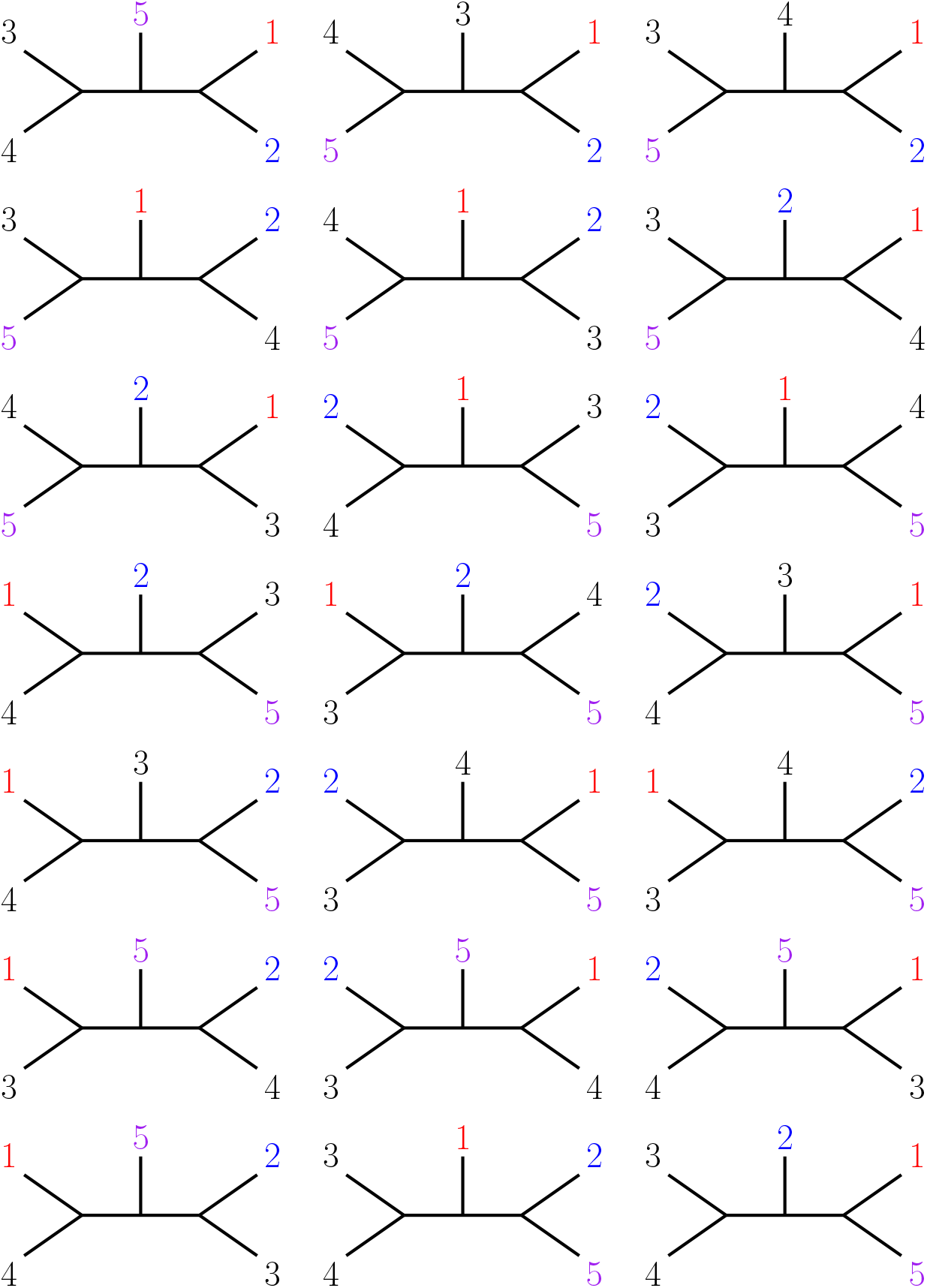
Type 2 labeled tree topologies for Property 1. These 21 labeled tree topologies correspond to Type-2 NJ cones (Section 3.1.1), where each equivalence class contains at least one, but not all, cherry-picking orders that violate Property 1. The labeled tree topologies are distinguished based on reflection symmetry with respect to the central taxon.

**Fig. B3:**
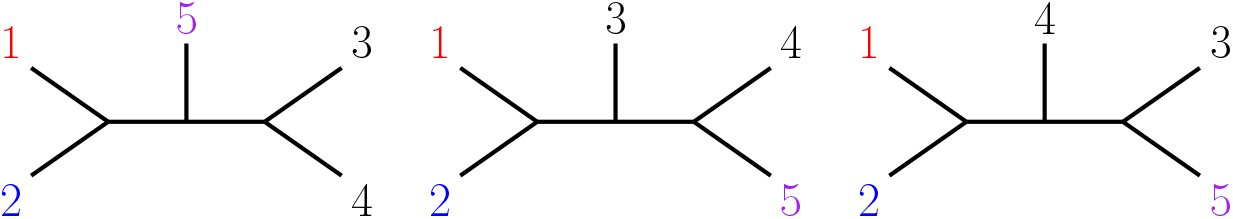
Type 3 labeled tree topologies for Property 1. These three labeled tree topologies correspond to Type-3 NJ cones (Section 3.1.1), where each equivalence class consists entirely of cherry-picking orders that violate Property 1. The labeled tree topologies are distinguished based on reflection symmetry with respect to the central taxon.

**Fig. B4:**
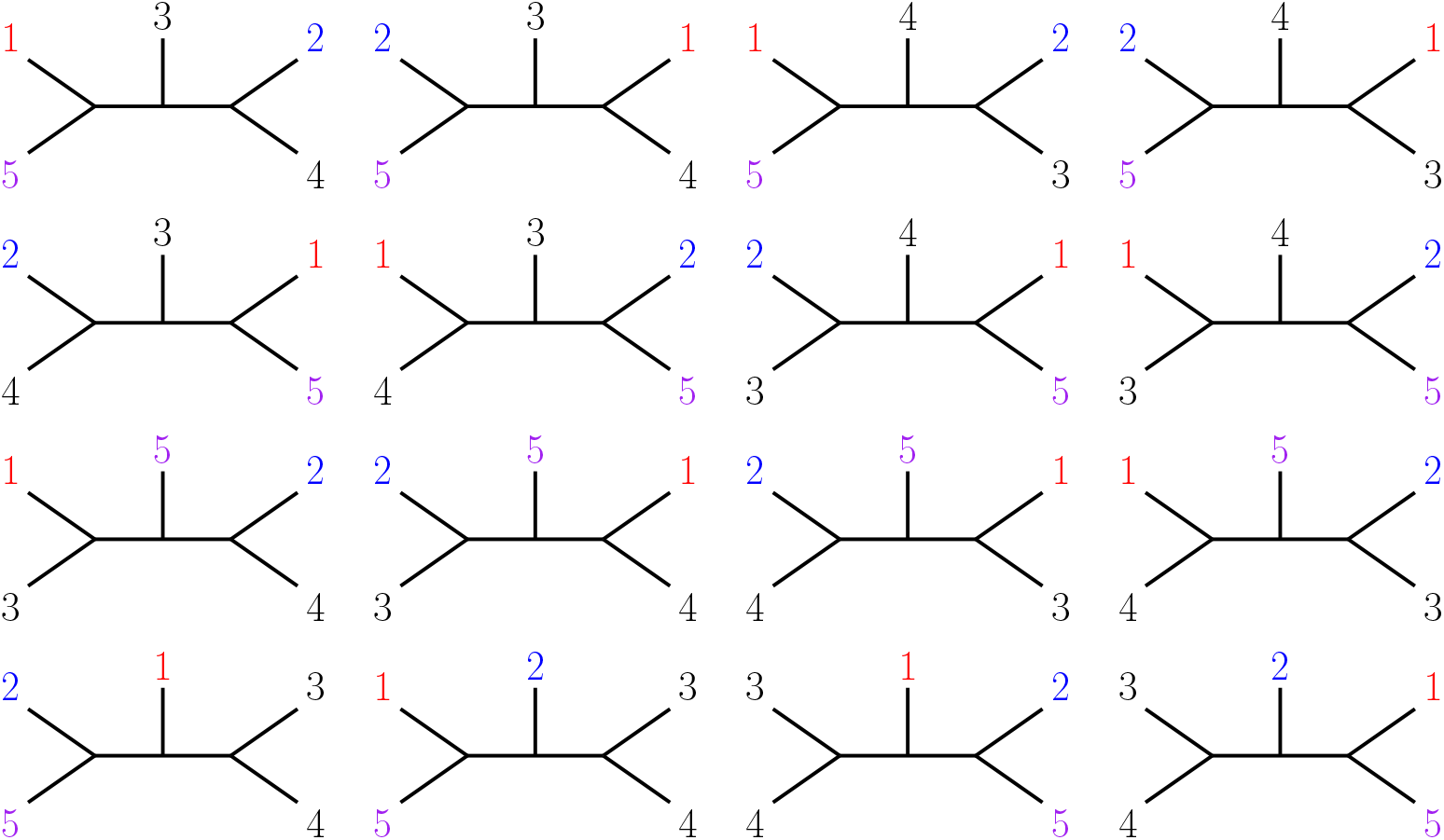
Labeled tree topologies that satisfy Property 3. These 16 labeled tree topologies represent the complete set of NJ cones that satisfy Property 3 (Section 3.1.2). The labeled tree topologies are distinguished by their reflection symmetry relative to the central taxon.

**Fig. B5:**
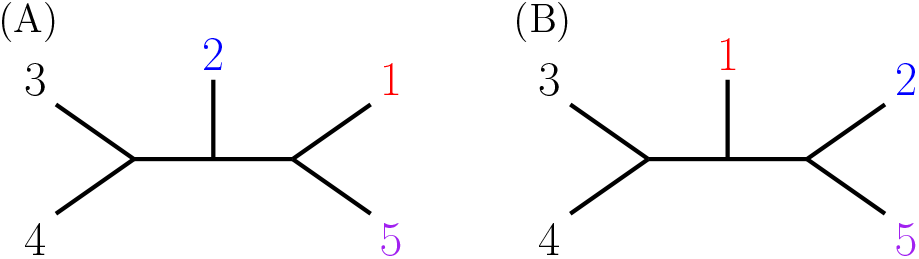
Two labeled tree topologies that always satisfy Property 2. For any admixture fraction *α* ∈ (0, 1), *π*_*α*_(*C*_(4,3)(5,1)_) and *π*_*α*_(*C*_(4,3)(5,2)_) are the only two induced NJ cones in which every dissimilarity vector d^(4)^ satisfies Property 2. **(A)** Labeled tree topology corresponding to the cone *C*_(4,3)(5,1)_. **(B)** Labeled tree topology corresponding to the cone *C*_(4,3)(5,2)_. The labeled tree topologies are distinguished based on reflection symmetry with respect to the central taxon.

